# Autophagosomes coordinate an AKAP11-dependent regulatory checkpoint that shapes neuronal PKA signaling

**DOI:** 10.1101/2024.08.06.606738

**Authors:** Ashley Segura-Roman, Y. Rose Citron, Myungsun Shin, Nicole Sindoni, Alex Maya-Romero, Simon Rapp, Claire Goul, Joseph D. Mancias, Roberto Zoncu

## Abstract

Protein Kinase A (PKA) is regulated spatially and temporally via scaffolding of its catalytic (Cα/β) and regulatory (RI/RII) subunits by the A-kinase-anchoring proteins (AKAP). PKA engages in poorly understood interactions with autophagy, a key degradation pathway for neuronal cell homeostasis, partly via its AKAP11 scaffold. Mutations in AKAP11 drive schizophrenia and bipolar disorders (SZ-BP) through unknown mechanisms. Through proteomic-based analysis of immunopurified lysosomes, we identify the Cα−RIα-AKAP11 holocomplex as a prominent autophagy-associated protein kinase complex. AKAP11 scaffolds Cα−RIα to the autophagic machinery via its LC3-interacting region (LIR), enabling both PKA regulation by upstream signals, and its autophagy-dependent degradation. We identify Ser83 on the RIα linker-hinge region as an AKAP11-dependent phospho-residue that modulates RIα-Cα binding and cAMP-induced PKA activation. Decoupling AKAP11-PKA from autophagy alters Ser83 phosphorylation, supporting an autophagy-dependent checkpoint for PKA signaling. Ablating AKAP11 in induced pluripotent stem cell-derived neurons reveals dysregulation of multiple pathways for neuronal homeostasis. Thus, the autophagosome is a novel platform that modulate PKA signaling, providing a possible mechanistic link to SZ/BP pathophysiology.

## Introduction

Master regulatory kinases such as Protein Kinase A (PKA), mechanistic Target of Rapamycin Complex 1 (mTORC1) and AMP-regulated Protein Kinase (AMPK) associate with different intracellular organelles such as mitochondria, lysosomes and lipid droplets, where they interact with specific upstream regulators and downstream substrates to carry out signaling responses that govern cell growth, differentiation, survival and metabolic adaptation ^1–5^.

PKA is a prominent Ser/Thr kinase that transduces cyclic adenosine monophosphate (cAMP) -mediated signals from many different hormones and neurotransmitters to instruct biological responses ranging from regulation of glucose and lipid metabolism in peripheral tissues, to memory formation and consolidation in the brain^6–9^. In its inactive state, PKA exists as a holoenzyme composed of a homodimer of two regulatory (R) subunits, each of which in turn holds and inhibits two catalytic subunits. Binding of cAMP to two nucleotide binding domains (NBD) on each R subunit induces a conformational change that frees up the C-subunit active site, enabling phosphorylation of a wide variety of downstream substrates^4,10,11^.

Four subtypes of R-subunits, RIα, RIβ, RIIα, and RIIβ, form as many distinct homodimers; in addition to C-subunits, each R-dimer, in turn, can bind to several A-kinase anchoring proteins (AKAPs). AKAP proteins organizes PKA signaling into a distributed intracellular network, where microdomains of PKA holoenzymes, scaffolded to different subcellular compartments, perform specific regulatory actions upon local elevation of cAMP levels^12^. Detailed mechanistic studies of AKAP proteins have conclusively demonstrated their ability to link PKA activity to specific organelles and cellular compartments, including the plasma membrane, nuclear envelope, mitochondria, Golgi apparatus, lipid droplets and centrosome^5,7^.

AKAP11 (also known as AKAP220) is among the least well understood AKAPs. This 220-KDa PKA-scaffolding protein has been implicated in processes such as actin turnover^13^, microtubule dynamics^14^, and osmoregulation^15^. AKAP11 has two binding sites for the regulatory subunits, one which binds to both RIα and RIIα, whereas the second one solely binds to RIIα^16^. Along with the PKA holoenzyme, AKAP11 scaffolds glycogen synthase kinase (GSK) IIIβ, possibly allowing cross-regulation between the two kinases^16,17^. AKAP11 also binds to protein phosphatase 1 (PP1), possibly as part of an integrated system to quickly activate and then shut down downstream substrates for both PKA and GSKIII-β^18^.

Recently, polymorphisms in the AKAP11 gene sequence that lead to protein truncation and loss-of-function have been strongly associated with increased risk for schizophrenia (SCZ) and bipolar disorder (BP) ^19–21^. A deep-proteomic study comparing brain tissue samples from SCZ and BP patients with synaptosomal preparations from AKAP11-knockout mice revealed shared alterations in the abundance of protein classes that are important for brain function, including mitochondria, vesicle trafficking and tethering^22^. However, the molecular links between AKAP11 deficiency and neuronal dysfunction that leads to SCZ and BP remain unknown.

Several reports have implicated AKAP11 in directing PKA to specific vesicle populations. AKAP11 was shown to recruit RIα to multivesicular bodies (MVBs), but only when RIα is not bound to Cα such as in high cAMP concentrations^23^. More recently, AKAP11 was proposed to function as a selective autophagy receptor that, by interacting with ATG8-family proteins, causes selective RIα capture and degradation into autophagosomes, while sparing Cα from degradation^24,25^.

Autophagy is a vesicular catabolic pathway that singles out proteins, macromolecules and organelles for capture within double-membraned autophagosomes, which subsequently deliver their cargo to lysosomes for degradation^26,27^. Recognition of autophagic substrates relies on a receptor-adaptor system^28,29^. Autophagic adaptors of the ubiquitin-like ATG8-family: MAP1LC3A/B/C, GABARAP and GABRAPBL1/2, are inserted in the autophagosomal membrane via their C-terminal conjugation to phosphatidylethanolamine (PE). ATG8-family adaptors recognize conserved linear motifs known as LC3-interacting regions (LIRs) on selective autophagic receptors, which in turn recruit various cargo to the nascent autophagosome^30,31^.

Consistent with the critical roles of autophagy in nutrient homeostasis and cellular quality control, this pathway is subjected to regulation by master nutrient-sensing kinases such as mTORC1, which negatively regulates the autophagosome-nucleating ULK1 complex under high nutrients, and AMPK, which stimulates several autophagic regulatory complexes during energy shortage^32,33^. Conversely, emerging evidence suggests that autophagy can exert feedback regulation on upstream signaling pathways: for example, nutrients generated by autophagic breakdown of macromolecules can reactivate mTORC1 and shut down AMPK signaling during recovery from starvation^33,34^. However, unlike lysosomes and mitochondria, which carry out well-recognized signaling functions on their cytoplasmic face that complement their internal (luminal) activities^33,35^, whether autophagosomes can act as organizers of cellular signaling at any point during their life cycle remains unclear.

The relationship between PKA signaling and autophagy is complex and not fully understood. Phosphoproteomic studies in yeast and human cells identified many autophagic regulators and effectors as PKA substrates, including the ATG1/ULK1 initiation complex, the ATG8 and ATG12 conjugation systems, as well as accessory factors and adaptors^36–39^. PKA-dependent phosphorylation of LC3B and ATG16L1 were shown to inhibit autophagosome formation in neurons and endothelial cells, respectively, suggesting that, like mTORC1, PKA may inhibit autophagy^40,41^. Conversely, functional genomic screens in yeast identified genes involved in each step of autophagy, from induction to substrate breakdown, as candidate negative regulators of PKA signaling^42^.

The findings that AKAP11 promotes autophagic degradation of RIα has led to a model in which selective RIα elimination increases the ratio of free C-subunit, thereby promoting PKA signaling^24,25^. This mechanism was proposed to potentiate C-mediated phosphorylation of PKA-cAMP response element-binding (CREB) transcription factor, boosting mitochondrial metabolism and conferring resistance to glucose deprivation^24^. Along similar lines, a further study proposed that autophagy-dependent elimination of RIβ in neurons frees up Cα to enhance PKA signaling at the synapse^43^.

However, these models need to be evaluated in light of well-established features of PKA regulation. First, the R-subunit does not play a purely inhibitory role toward the C-subunit. On the contrary, by scaffolding C to different AKAP proteins, R helps place the former in proximity to numerous organelle-specific substrates^5,7,44,45^. Thus, decreasing the stoichiometry of R-to C-subunits would be expected to have mixed effects on downstream signaling, favoring some phosphorylation events while decreasing others. Second, recent evidence suggests that, at physiological cAMP levels found within the cell, C-subunits do not completely separate from the R-subunit-AKAP complex. Instead, the three components remain bound in an open, signaling-competent conformation that is capable of substrate recognition and phosphorylation^46,47^. Thus, autophagic capture of AKAP11-RIα seems unlikely to completely spare C-subunits from degradation.

To achieve a fine-grained understanding of how autophagy regulates PKA, and possibly other multiprotein signaling complexes, we combined immunoisolation and proteomic-based profiling of lysosomes from autophagy competent and deficient cells with bioinformatic analysis of protein-protein interaction datasets. These experiments identify the PKA holoenzyme, primarily composed of the RIα and Cα subunits and AKAP11, as a kinase complex prominently associated with autophagy. In agreement with previous reports, AKAP11 bridges RIα to LC3B and GABARAP via its LIR motif. However, we find that Cα is also scaffolded to LC3B and GABARAP via its association with RIα and AKAP11. Crucially, we find that association with the autophagic machinery modulates PKA-dependent signaling, both by co-scaffolding of the PKA holoenzyme with upstream regulators, and by promoting its degradation. In particular, phosphoproteomic analysis in iPSC-derived neurons reveals AKAP11-dependent RIα phosphorylation on a critical regulatory site; accordingly, depleting AKAP11 or disabling its interaction with the autophagy machinery led to dysregulation of numerous downstream signaling programs linked to neuronal cell homeostasis. Collectively, our data point to the autophagosome as a central signaling hub that plays a critical role in shaping PKA-dependent signaling responses via AKAP11. Alteration of these signaling programs upon AKAP11 loss may contribute to the pathophysiology of SZ/BP.

## Results

### Proteomic and bioinformatic identification of multiprotein autophagic substrates

We reasoned that signaling complexes that interact tightly with the autophagic machinery may be subjected, at least in part, to autophagy-dependent degradation. To identify these autophagy-associated signaling complexes, we thus carried out immunoisolation of lysosomes (the endpoint of autophagy) followed by their proteomic-based profiling from control HEK-293T cells versus HEK-293T cells deleted for an essential autophagic regulator, the E1-like factor ATG7 ^48,49^ (Fig. 1A and Fig. S1A-S1B). To unambiguously distinguish lysosomal resident proteins from *bona fide* substrates, we compared lysosomal immunoprecipitates from DMSO-treated control and ATG7-KO cells, versus cells treated with the protease inhibitor cocktail, leupeptin + pepstatin (L+P)^48^. Proteins that showed selective accumulation in L+P-treated control lysosomes included canonical autophagic receptors CALCOCO2, NBR1, TAX1BP1 and NCOA4, along with transmembrane and extracellular proteins that reach the lysosome via endocytic uptake and endosomal trafficking (Supplementary data 1). As expected, autophagic receptors were depleted in lysosomes immunopurified from ATG7-KO cells (Supplementary data 1), establishing a clear pattern for recognition of *bona fide* autophagic substrates.

**Figure 1.**
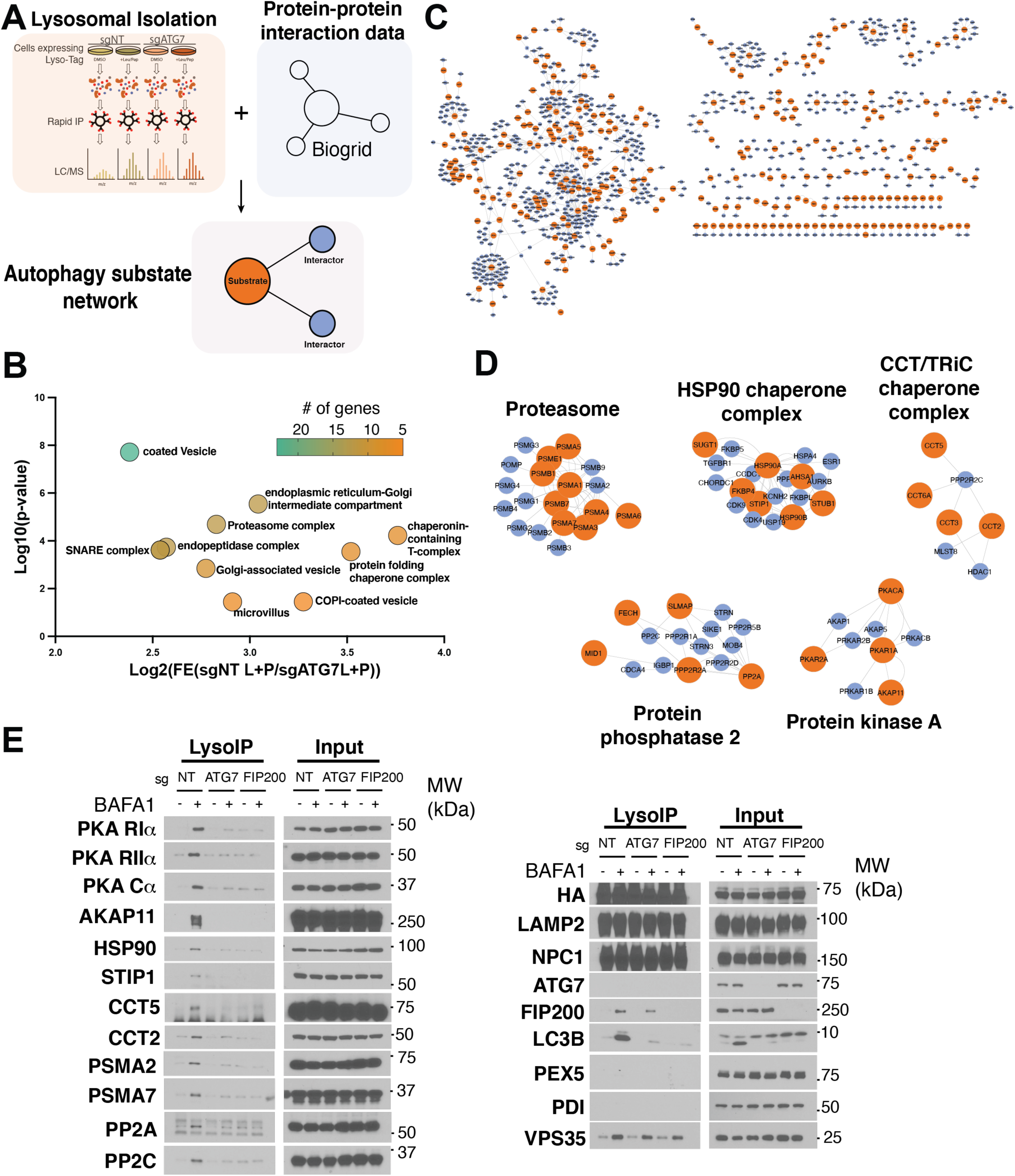
Multiprotein complexes are autophagic substrates. **A**. Summary chart of the workflow for identification of autophagy-dependent degradation of multi-protein complexes. HEK293T sgNT or sgATG7 cells were treated with 20*µ*M Leupeptin + 20*µ*M Pepstatin or DMSO for 24 hours prior to lysosomal immunoprecipitation. Resulting proteomic analysis from proteins classified as autophagic-dependent lysosomal substrates were integrated into a network using protein-protein interaction data from Biogrid database. 4 citations were required for protein-protein interaction. **B.** Plot of “cellular component slim” GO-terms enriched in wild-type lysosomes compared to autophagy-null lysosomes in the Leupeptin + Pepstatin condition. **C.** Network representation of autophagy-dependent substrates (orange). Blue nodes show cited protein-protein interactions to provide context for which multi-protein complex substrates belong to. **D.** Highlighted multi-protein complexes from protein network analysis. Proteins belonging to the proteasome, HSP90 Chaperone complex, CCT/TRiC chaperone complex, protein phosphatase 2, or protein kinase A complexes are enriched as autophagy-dependent lysosomal substrates. **E** Immunoblots of lysosomal immunoprecipitation and corresponding input from HEK293T sgNT, sgATG7 or sgFIP200 after treatment with 500nM bafilomycin A1 (BafA1) for 5 hours to block lysosomal degradation of autophagic substrates.

Gene Ontology (GO) analysis showed differential accumulation of several classes of putative autophagic substrates in lysosomes from control, L+P treated cells versus ATG7-KO, L+P-treated cells. The most autophagy-dependent substrate categories included factors involved in protein folding, transport and quality control (Fig. 1B and Fig. S1C).

We next analyzed the proteomic data through a custom-written bioinformatic pipeline that clusters proteins based on physical interaction (BioGRID database) (Fig. 1A and S1B). The resulting network view identified several multiprotein complexes as novel substrates of autophagic degradation (Fig. 1C). We identified complexes involved in protein quality control, including the proteasome (12 subunits detected with p-value =2.05E-05) and the CCT/TRiC chaperone complex (7 subunits) (Fig. 1D and Supplementary data 1). Also detected was the HSP90 chaperone complex, including HSP90α class A and B, their p23/PTGES3 and FKBP4 cofactors, as well as HSP70, which, while not stably bound to HSP90, functions in close association with HSP90 on substrate proteins^50^ (Fig. 1D and Supplementary data 1). The enrichment of protein quality control factors likely reflects their abundant levels in the cell, as well as a possible high rate of folding and assembly failure of their client proteins under steady-state conditions.

Our pipeline also identified multiprotein complexes involved in signal transduction, specifically the catalytic and one regulatory subunits of protein phosphatase 2 (PPP2CA and PPP2R1A, respectively), and the PKA enzyme complex composed of the Cα catalytic subunit, the RIα and RIIα regulatory subunits, and the AKAP11 (also known as AKAP220) scaffolding protein (Fig. 1D and Supplementary data 1).

### The PKA holoenzyme is a substrate for autophagic degradation

To independently validate the proteomic data, we immunoisolated lysosomal samples from control and ATG7-KO HEK-293T cells, as well as from cells deleted for FIP200, a component of the ULK1 complex that is essential for autophagy initiation^26^. We carried out lyso-IP both in baseline conditions and following treatment with the vacuolar H^+^ATPase (v-ATPase) inhibitor, Bafilomycin A1 (BafA1), which, like L+P, inhibits substrate breakdown within the lysosomal lumen (Fig. 1E). Immunoblotting of these samples confirmed the autophagy-dependent degradation of the multiprotein complex components detected by mass spectrometry, including the proteasome, HSP90, CCT/TriC, PP2A (Fig. 1E). Supporting autophagic degradation of the PKA holoenzyme, RIα, RIIα, Cα and AKAP11 were clearly detected in lysosomes from control cells treated with BafA1, but not in lysosomes from BafA1-treated ATG7- and FIP200-KO cells (Fig. 1E). In light of recently proposed models of autophagy-dependent regulation of PKA^24,25,43^, and the recently discovered association of AKAP11 mutations with SCZ and BP^19–21^, we decided to further investigate the Cα-R1α-AKAP11 complex as a potential autophagy interactor and substrate.

To gain a quantitative understanding of PKA-AKAP11 capture by autophagy, we visualized the lysosomal mass spectrometry data (Supplementary data 1) as volcano plots of Log2 fold change over -Log10 p-value. These plots showed that RIα is among the most enriched proteins by L+P treatment, on par with canonical autophagic receptors such as TAX1BP1 and p62, whereas Cα and AKAP11 were stabilized to a lesser degree (Fig. 2A). The same relative enrichment was found when comparing lysosomes from L+P-treated wild-type versus ATG7-deleted cells (Fig. 2B).

**Figure 2.**
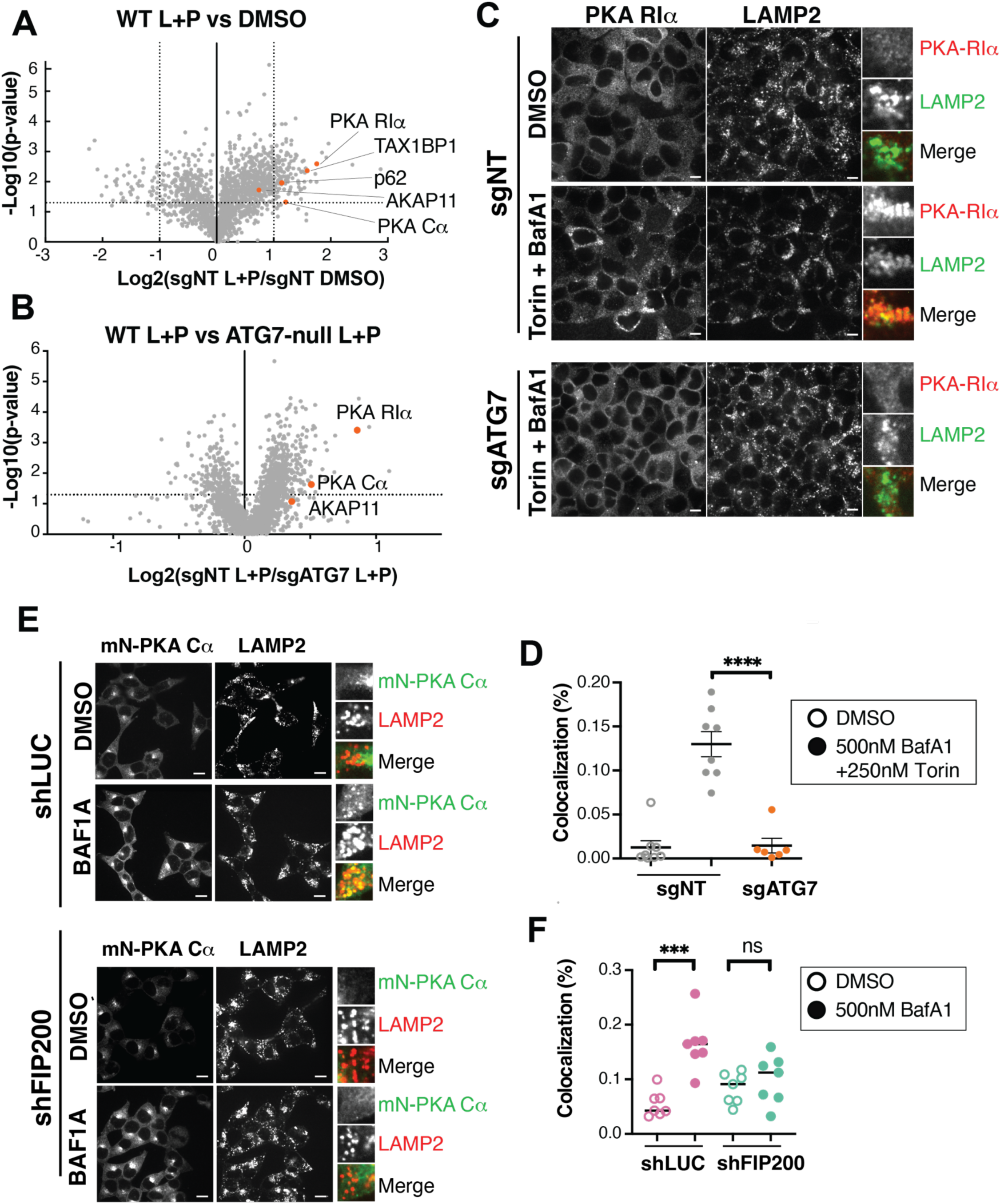
The PKA holoenzyme is degraded via autophagy. **A, B.** Proteomic analysis of Lyso-immunoprecipitation (LysoIP) from HEK293T sgNT and sgATG7. (A) Volcano plots show the ratio of 20*µ*M Leupeptin + 20*µ*M Pepstatin (L+P) to DMSO treated for sgNT. (B) Volcano plots shows the ratio of sgNT L+P treated to sgATG7 L+P treated. PKA signaling complex highlighted in orange circles, along with canonical autophagic receptors. **C.** Immunofluorescence of HEK293T sgNT and sgATG7 that over-express Flag-PKA RIα were treated with 5-hou 500nM BafA1 and 250nM Torin for 1 hour before fixation and immunostaining for Flag and LAMP2. 10μm scale bar. **D.** Quantification of Flag and LAMP2 co-localization from 8 non-overlapping fields. ****p(adj)<0.0001. **E.** Immunofluorescence of HEK293T endogenously expressing mNeonGreen-PKA Cα in cells treated with shLuciferase control or shFIP200. Cells were serum-starved O/N then treated with 500nM BafA1 or DMSO for 5 hours before fixing and immunostaining for mNEONGREEN and LAMP2. 10μm scale bar **F.** Quantification of the co-localization of mNeonGreen-PKA Cα with LAMP2 under all conditions for 8 non-overlapping fields. ***p(adj)<0.0002.

Autophagy-dependent capture of the PKA holoenzyme was also evident in immunofluorescence-based experiments. In cells stably expressing FLAG-tagged RIα, blocking lysosomal proteolysis with BafA1 led to pronounced accumulation of FLAG-R1α in LAMP2-positive lysosomes of control, but not ATG7-deleted, HEK-293T cells (Fig. 2C and 2D). Similarly, in HEK-293T cells in which endogenous Cα is C-terminally tagged with mNeon^51^, treatment with BafA1 led to pronounced accumulation of Cα−mNeon in LAMP2-positive lysosomes that was abolished by shRNA-mediated knock down of FIP200, consistent with autophagy-dependent capture and degradation of Cα (Fig. 2E and 2F and Fig. S1D).

In conclusion, converging evidence from organelle proteomics and immunolocalization support autophagic capture and lysosomal degradation of both the regulatory and catalytic PKA subunits via an ATG7- and FIP200-dependent mechanism.

### AKAP11 mediates capture of the PKA holoenzyme

It was previously reported that AKAP11 promotes autophagic degradation of a fraction of the RIα pool^24,25^. However, our detection of Cα in immunopurified lysosomes, as well as by colocalization analysis in intact cells, indicate that the autophagic machinery interacts with not just RIα, but with the PKA holoenzyme (Fig. 1D-1E, 2E-2F). To verify that autophagic capture of Cα is also AKAP11-dependent, we stably expressed the lysosomal affinity tag in both control and AKAP11-deleted HEK-293T cells, and immunoblotted lysosomal samples for multiple PKA proteins. This analysis clearly showed that both Cα and RIα were strongly depleted from lysosomes immunopurified from AKAP11-KO cells (Fig. 3A).

**Figure 3.**
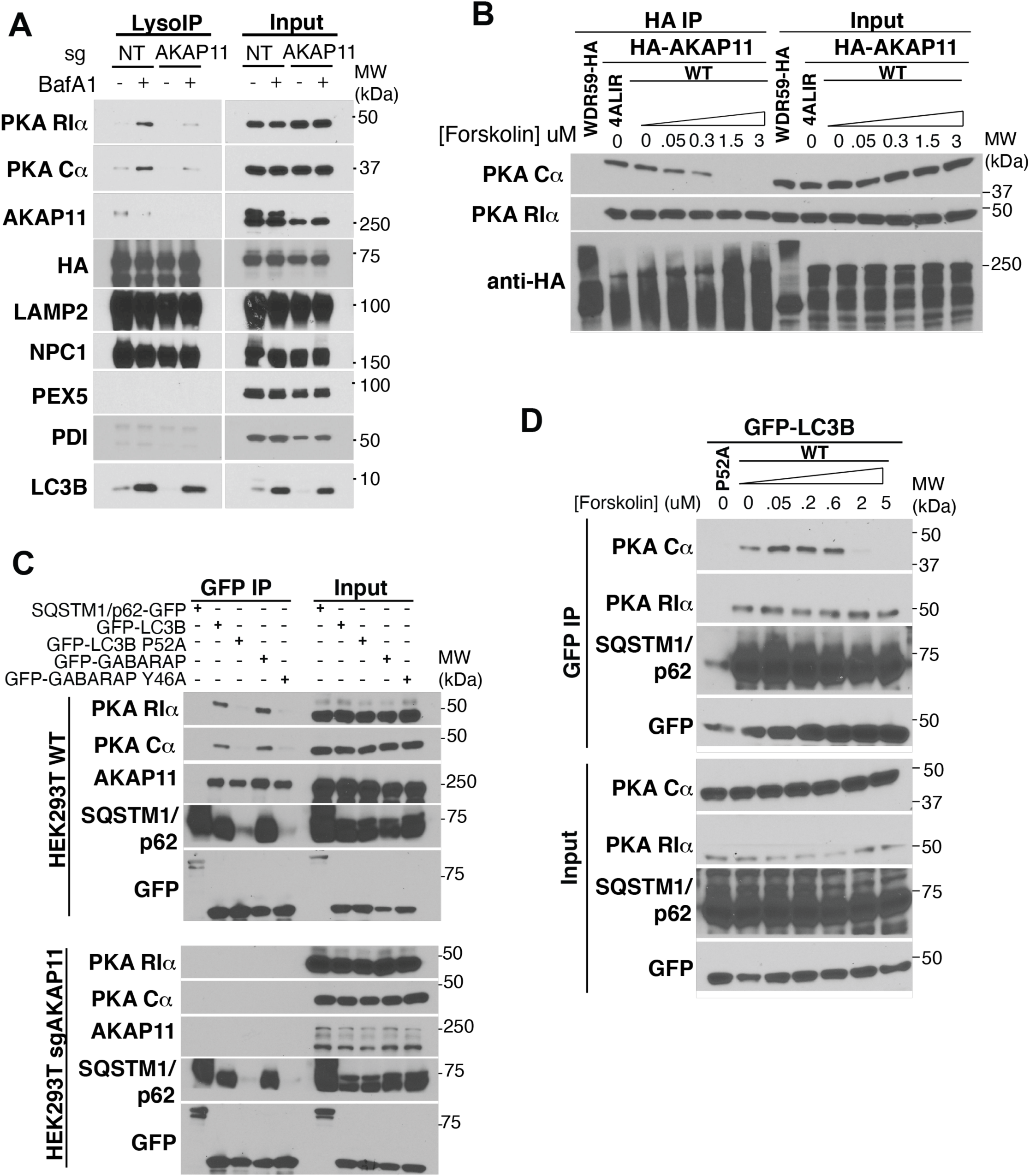
The PKA holoenzyme is degraded via AKAP11. **A.** Immunoblots of LysoIP samples from HEK293T sgNT or sgAKAP11 cells, treated with 500nM BafA1 or DMSO for 5 hours. **B.** Immunoblots from HEK293T which were transiently transfected with the WDR59-HA or indicated HA-AKAP11 cDNA constructs (the 4ALIR mutant is described in Fig. 4). Cells were treated with increasing concentrations of Forskolin, as indicated, for 20 minutes in complete media (10% FBS DMEM). **C**. Immunoblots from GFP immunoprecipitation from HEK293T sgNT or sgAKAP11 cells stably expressing the indicated GFP-tagged protein. **D.** Immunoblots from HEK293T sgNT and sgAKAP11 cells stably expressing GFP-LC3 WT or GFP-LC3 P52A mutant. Cells were treated with increasing concentrations of Forskolin, as indicated, for 20 minutes in complete media (10% FBS DMEM) before lysis and GFP immunoprecipitation

Whereas the R−AKAP interaction is relatively stable (in the nanomolar range), Cα progressively dissociates from the R-AKAP complexes as cytoplasmic concentrations of cAMP increase^52^. However, recent structural and biochemical evidence indicates that concentrations of cAMP that are sufficient to activate PKA signaling do not cause complete Cα dissociation from the R-AKAP complex. Instead, the AKAP-holoenzyme remains associated while adopting a range of dynamic conformations that are compatible with substrate engagement and phosphorylation^46,47^. In line with this idea, HA-tagged AKAP11 co-immunoprecipitated both RIα and Cα at the steady-state levels of cAMP that exist under full media conditions. To achieve complete separation of Cα from the R1α−AKAP11 complex, we had to treat cells with the adenylyl cyclase activator, forskolin, at >1μM concentrations, which lead to supra-physiological cAMP levels in the cell^47^ (Fig. 3B).

To be captured into autophagosomes, substrate proteins must interact with Atg8-family proteins LC3A/B/C or GABARAP, either via an LC3-interacting region (LIR) within their amino acid sequence, or indirectly through a selective autophagic receptor^31,53^. In co-immunoprecipitation (co-IP) experiments, GFP-tagged LC3B and GABARAP bound to R1α, Cα and AKAP11 with relatively high strength (Fig. 3C). The RIα and Cα interactions were largely abolished when using point mutants in the hydrophobic groove of LC3 and GABARAP that impair recognition of canonical LIR motifs (P52A and Y46A, respectively)^30,31^ (Fig. 3C, top). Supporting the requirement for AKAP11 in autophagic capture of the PKA holoenzyme, GFP-LC3B and GFP-GABARAP failed to immunoprecipitated endogenous RIα and Cα when expressed in AKAP11-deleted cells, whereas their binding to p62/SQSTM1 was unaffected by lack of AKAP11 protein (Fig. 3C, bottom).

Similar to HA-AKAP11, GFP-LC3B co-immunoprecipitated both RIα and Cα in full media conditions. Stimulating GFP-LC3B-expressing cells with increasing concentrations of forskolin progressively reduced the amount of Cα immunoprecipitated by GFP-LC3B, consistent with Cα interacting with LC3B indirectly, via its association with R1α (Fig. 3D). However, as for the AKAP11 co-IP experiments (Fig. 3B), Cα fully separated from LC3B only at supraphysiological concentrations of forskolin (>1μM) (Fig. 3D).

Thus, under steady-state conditions autophagy captures and degrades the AKAP11-bound pool of PKA holoenzyme via association of both RIα and Cα with AKAP11 and LC3/GABARAP.

### AKAP11 binding to LC3 and R1α is required for autophagic degradation of the PKA holoenzyme

We next tested the mechanisms underlying autophagic capture of the AKAP11-containing PKA holoenzyme. Human AKAP11 contains a putative LIR motif, WSNL, at a.a. 1736-1739^24^. Wild-type, V5-tagged AKAP11 (stably expressed in AKAp11-deleted HEK-293T cells) was readily detected within LAMP2-positive lysosomes upon BafA1-treatment; in contrast, mutating the WSNL motif to four alanines caused the resulting AKAP11^4αLIR^ mutant to be largely excluded from lysosomes and instead maintain a diffuse staining pattern upon BafA1 treatment (Fig. 4A-4B).

**Figure 4.**
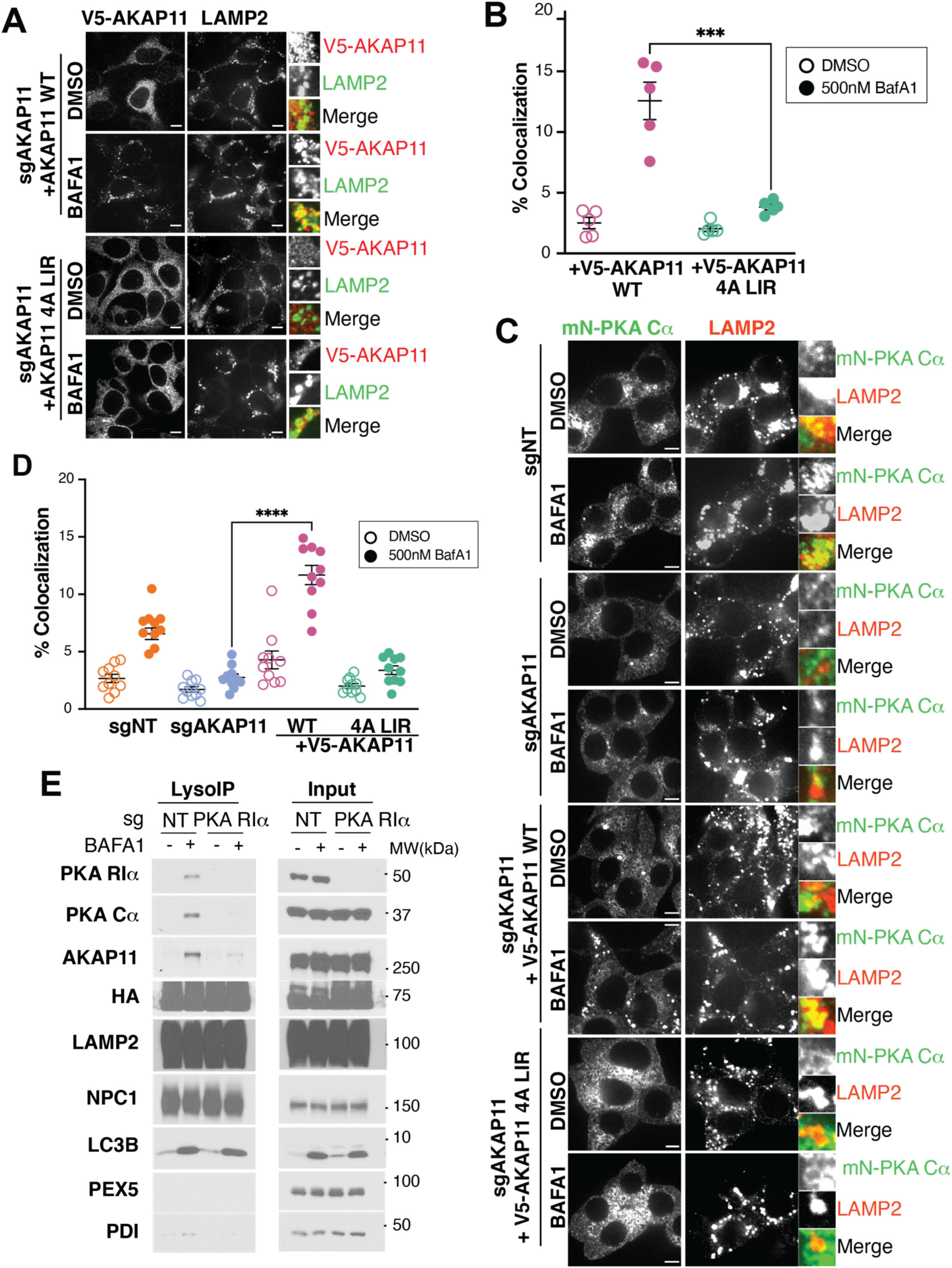
The AKAP11 LiR motif mediates autophagic capture of the PKA holoenzyme. **A.** Immunofluorescence from HEK293T sgAKAP11 cells stably expressing V5-AKAP11 WT or V5-AKAP11 4ALIR (WSNL>AAAA) were treated with 500nM BafA1 or DMSO for 5 hours before fixing and immunostaining for V5 and LAMP2. 10μm scale bar. **B.** Quantification of V5 and LAMP2 colocalization from 6 non-overlapping fields; ***p(adj.)=0.0005, unpaired t test. **C.** Immunofluorescence of HEK293T sgNT or sgAKAP11 with endogenously-tagged mNEONGreen-PKA Cα also stably expressing V5-AKAP11 construct as indicated. Cells were serum-starved O/N then treated with 500nM BafA1 or DMSO for 5 hours before immunostaining with LAMP2. **D.** Quantification of mNEONGreen and LAMP2 colocalization from 10 non-overlapping fields. ****p(adj)<0.0001, unpaired t test. **E.** Immunoblots of LysoIP samples from HEK293T sgNT or sgPKA RIα, treated with 500nM BafA1 or DMSO for 5 hours.

As the 1736-1739 LIR motif of AKAP11 was shown to be required for AKAP11-mediated R1α degradation^24^, we sought to determine its requirement for autophagic capture of Cα as well. For this, we ablated AKAP11 in HEK-293T cells in which endogenous Cα is tagged with mNeon, and reconstituted them with stably expressed wild-type AKAP11 or with the AKAP11^4ALIR^ mutant. AKAP11^4ALIR^ bound to endogenous Cα and RIα with identical strength to the wild-type protein (Fig. 3B).

Immunofluorescence analysis showed that ablating AKAP11 blocked lysosomal localization of Cα-mNeon in BafA1-treated cells (Fig. 4C-4D). Cα-mNeon – LAMP2 colocalization was restored by re-expressing AKAP11^WT^, but not by the AKAP11^4ALIR^ mutant (Fig. 4C-4D). Thus, while dispensable for the integrity of the Cα− RIα-AKAP11 complex, the LIR motif of AKAP11 appears critical for autophagic capture of the PKA holoenzyme.

Because RIα bridges AKAP11 with Cα (Fig. 3B), we also examined potential roles for RIα in promoting autophagic capture of the holoenzyme. We generated RIα-deleted HEK-293T cells via CRISPR-Cas9, and carried out lysosome affinity purification and immunoblotting from these cells. As expected, deleting RIα prevented accumulation of Cα in lysosomes (Fig. 4E). Interestingly, the levels of lysosomal AKAP11 were also suppressed in lysosomes from RIα−deleted cells, suggesting that AKAP11 must be bound to RIα in order for its capture by the autophagic machinery to occur (Fig. 4E).

Together, these data show that AKAP11 and RIα cooperatively scaffold the entire Cα− RIα-AKAP11 complex to autophagosomes via LIR-dependent interactions between AKAP11 and the autophagic machinery.

### Phosphoproteomic analysis in iNeurons reveals AKAP11-dependent PKA regulation

The ability of AKAP11 to both scaffold its bound PKA holoenzyme to the autophagosome, and to promote its autophagy-dependent degradation, prompted us to dissect its regulatory roles toward PKA signaling. Because AKAP11 is highly expressed in the brain, and given the strong association between AKAP11 gene truncations and SZ/BP^19–21^, we carried out these experiments in induced pluripotent stem cell (iPSC)-derived neurons. Using CRISPR-interference (CRISPRi), we first suppressed AKAP11 expression in iPSCs that can be differentiated into inducible cortical neurons (i3 neurons) via doxycycline-driven expression of neurogenin-2 (NGN2)^54^. Both prior to differentiation and at 7 days post-NGN2 induction, we verified efficient silencing of AKAP11 by three independent single-guide RNAs (sgRNAs). Consistent with the results in HEK-293T cells, all AKAP11-targeting sgRNAs led to strong stabilization of RIα and, to a lesser extent, of Cα (Fig. 5A, and Fig. S2A-S2D). AKAP11-depleted i3 neurons were morphologically similar to control neurons (Fig. 5B).

**Figure 5.**
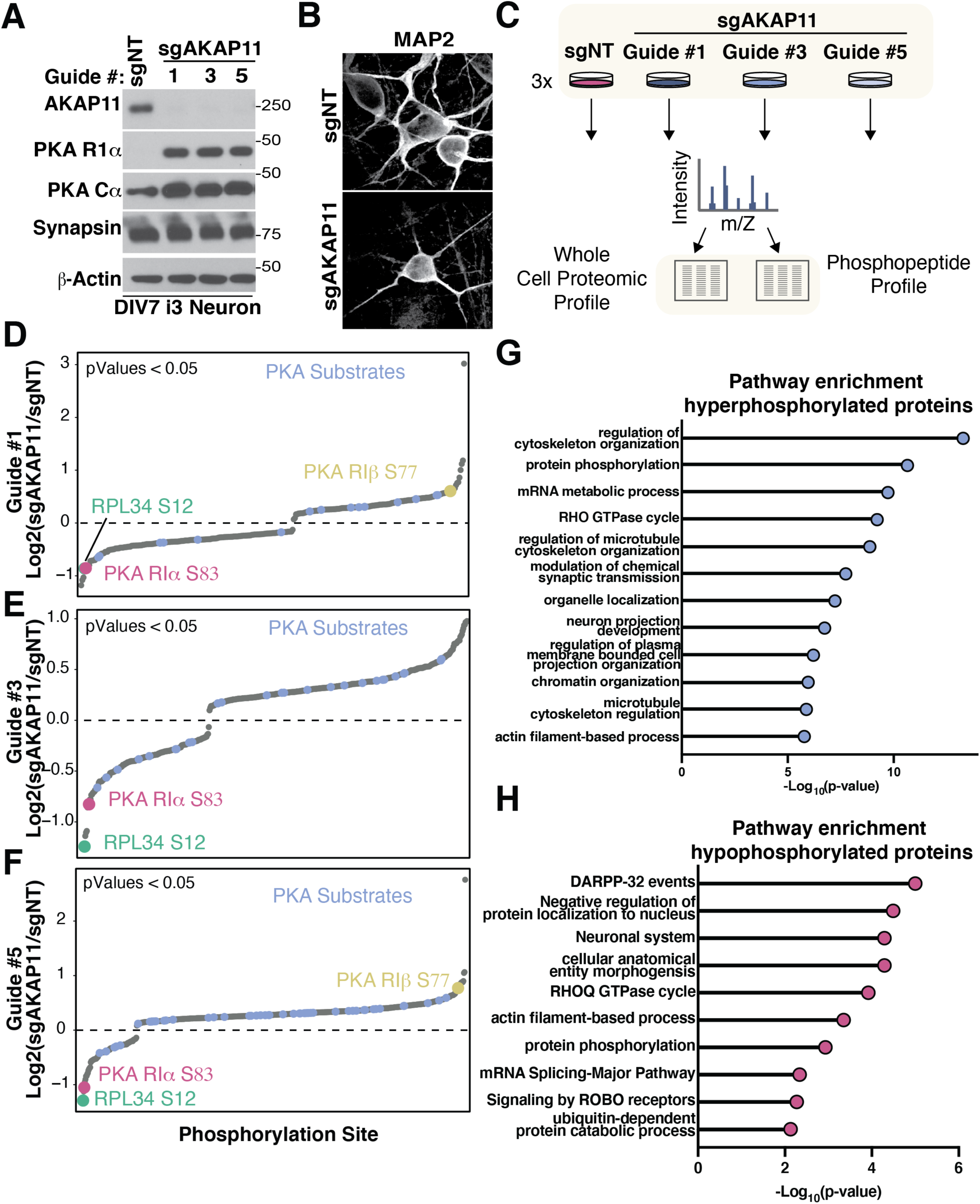
AKAP11 regulates PKA-dependent and independent signaling in i3 Neurons. **A.** Immunoblot of sgNT or sgAKAP11 i3 Neurons (DIV7) depicting degree of AKAP11 KD in the indicated guide. **B.** Immunofluorescence images of sgNT or sgAKAP11 i3 Neurons (DIV7) immunostained with MAP2. **C.** Schematic depicting mass spectroscopy workflow. Mass spectroscopy analysis of phosphorylated peptides was performed comparing i3 Neurons with three sgRNA guides (#1,3,5) against AKAP11 vs a non-targeting guide. **D, E, F.** After normalizing each phosphorylated peptide to the total protein abundance in the sample, waterfall plots were generated of the foldchanges with statistical significance (p-value < 0.05) comparing each of the three AKAP11 guides to the non-targeting guide. **G, H.** Bar graph depicting pathway enrichment analysis of phosphoproteomic data set. (G) pathway enrichment analysis of phosphoproteins enriched in sgNT i3 Neurons. Phosphorylated proteins with Log_2_Foldchange ≥ 0.2 and significant p-value (<0.05) were considered for pathway enrichment analysis. (H) pathway enrichment analysis of phosphoproteins depleted in sgAKAP11 i3 Neurons. Phosphorylated proteins with Log_2_Foldchange of ≤-0.2 and significant p-value (<0.05) were considered for pathway enrichment analysis.

We employed phosphoproteomic analysis to determine the effect of AKAP11 loss on PKA-dependent signaling in i3 neurons (Fig. 5C). Because RIα and Cα stabilization was clearly detectable under standard growth conditions (Fig. 5A and S2B-S2D), we carried out our analysis without further stimulating PKA activity or autophagy. We detected 228 peptides, belonging to 174 proteins, phosphorylation of which was significantly (p<0.05) altered in at least 2 out of 3 sgAKAP11 i3neuron lines (Fig. 5D-5F and supplementary data 2: phospho-peptide signals normalized by total protein levels). Notably, only 5.7% (13 sites) of differentially phosphorylated peptides matched the consensus PKA target site, whereas 94.3% were not predicted to be PKA sites (supplementary data 2).

Pathway enrichment analysis of differentially phosphorylated proteins revealed that, in line with its reported functions in non-neuronal lines^8^/6/24 3:23:00 PM AKAP11 controls phosphorylation of proteins involved in microtubule polymerization/depolymerization and regulation of the actin cytoskeleton^13–15^ (Fig. 5G-5H and supplementary data 2). Other differentially phosphorylated categories included synaptic function, neuronal morphogenesis and signaling (Fig. 5G-5H and supplementary data 2).

Of the 13 *bona fide* PKA target sites differentially regulated in an AKAP11-dependent manner, 7 sites were hypophosphorylated in AKAP11-depleted neurons, whereas 6 sites were hyperphosphorylated. A notable site was Ser12 on the large ribosomal protein 34 (RPL34), which was among the most hypophosphorylated sites in all three AKAP11-depleted lines (Fig. 5D-5F and supplementary data 2). Ser12 is highly conserved, and is located in the extended N-terminal region of RPL34, and lies in close proximity to several ribosomal RNA (rRNA) molecules within the large subunit core^55^ (Figure S3A-S3B). The position of Ser12 deep within the large ribosomal subunit core predicts that its PKA- and AKAP11-dependent phosphorylation could significantly impact ribosomal large subunit assembly and stability. According to published phosphoproteomic datasets (www.phosphosite.org), Ser12 is the most commonly detected phospho-site on RPL34, supporting its role as a putative regulatory site for the ribosome.

Among the most hypo-phosphorylated non-PKA substrate peptides in all three AKAP11-deleted i3 neuron lines was Ser83 in the RIα subunit (Fig. 5D-5F and supplementary data 2). Ser83 is highly conserved, is located within the linker–hinge region of R1α, N-terminally proximal to the inhibitory sequence (IS) (Figure S3C-S3D), and phosphorylation of several residues in this region had previously been proposed to regulate the stability of IS binding to the catalytic cleft of Cα and, thus, the efficiency of Cα inhibition^56,57^. Ser83 is a commonly detected phospho-site on RIα^57–59^ (www.phosphosite.org), thus it is likely to represent a *bona fide* regulatory site that, upon phosphorylation by upstream kinases (see Discussion) may fine-tune the activity levels of AKAP11-bound PKA.

Finally, in addition to RIα, its brain-specific paralog RIβ, and Cα levels, proteomic analysis revealed significant changes in the total levels of numerous other proteins (Fig. S2B-S2D and supplementary data 2). Protein classes that were especially increased in AKAP11-depleted i3 neurons included ribosome biogenesis and rRNA processing factors, an effect possibly linked to our finding of AKAP11- and PKA-dependent RPL34 phosphorylation on Ser12.

In summary, our phosphoproteomic analysis in i3 Neurons reveals a complex network downstream of AKAP11, whereby both PKA-dependent and PKA-independent phosphorylation events are either promoted or antagonized in an AKAP11-dependent manner.

### AKAP11-dependent RIα phosphorylation at Ser83 modulates PKA activation

The phosphoproteomic results in AKAP11-depleted i3 neurons are consistent with two possible roles for the interaction of the Cα−RIα-AKAP11 complex with the autophagic machinery. One is a scaffolding function that facilitates PKA-dependent phosphorylation of a subset of downstream substrates, as well as regulation of PKA itself by upstream regulators; the second is an inhibitory role, in which autophagy-dependent degradation of the PKA holoenzyme suppresses phosphorylation of other PKA substrates.

To dissect how these activating and inhibitory functions relate to one another, we carried out parallel whole-cell phosphoproteomic analysis in control, AKAP11-deleted and AKAP11-deleted cells reconstituted with exogenous wild-type AKAP11 or with the AKAP11^4ALIR^ mutant (Fig. 6A). We employed HEK-293T cells for these experiments as they were more amenable than the i3 neuron system to deletion-reconstitution protocols. As expected, re-expressing (V5-tagged) wild-type AKAP11 in sgAKAP11 cells decreased the elevated endogenous RIα levels, whereas V5-AKAP11^4ALIR^ failed to bring RIα down to wild-type levels, albeit not completely (Fig. S4A).

**Figure 6.**
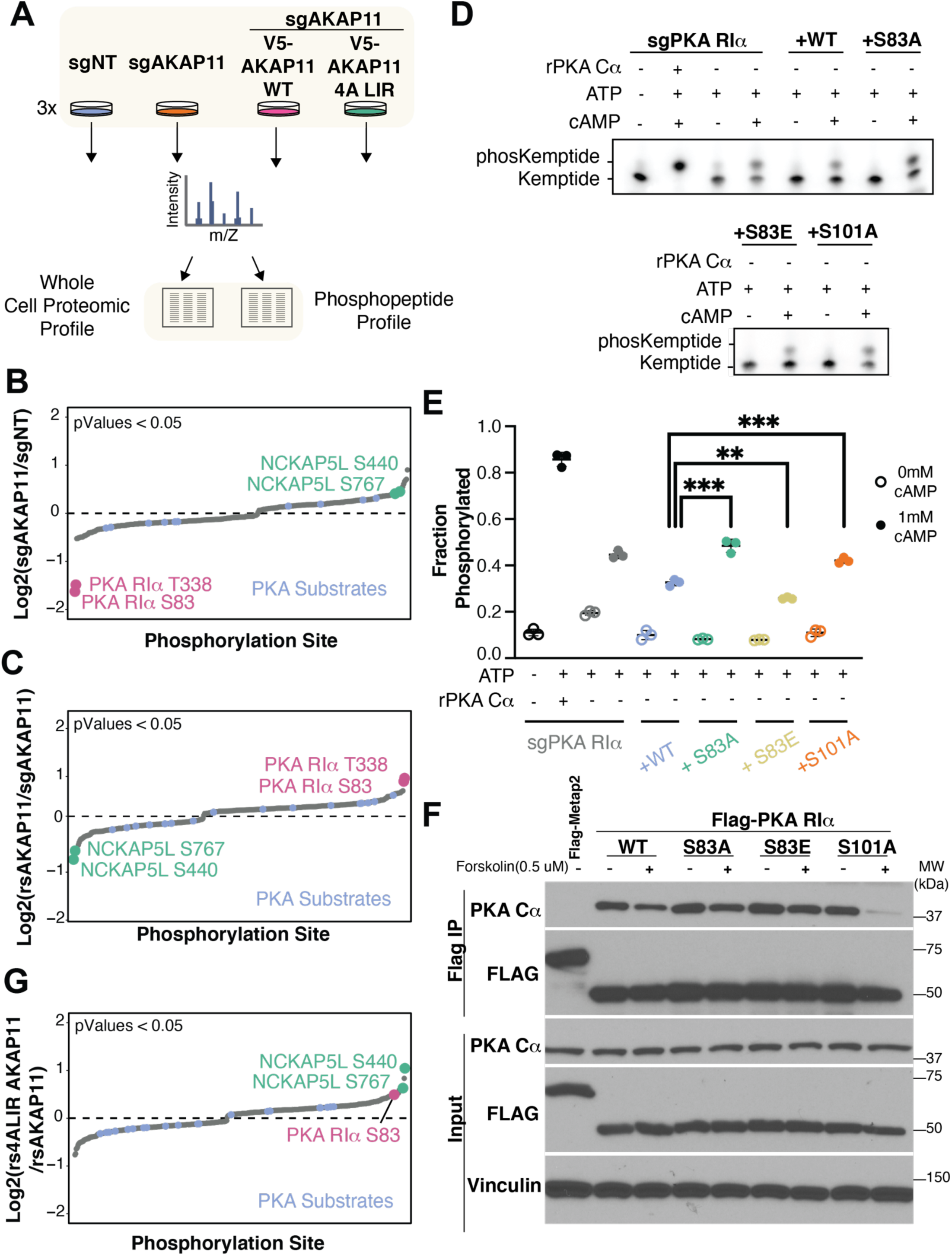
AKAP11 Regulates phosphorylation of RIα to modulate PKA activity. **A.** Schematic depicting mass spectroscopy workflow. **B, C.** Mass spectroscopy analysis of phosphorylated peptides was performed comparing AKAP11 genotypes. After normalizing each phosphorylated peptide to the total protein abundance in the sample, waterfall plots were generated of the foldchanges with statistical significance (p-value < 0.05) comparing (B) sgAKAP11 knockout cells to control sgNT cell and (C) rescue V5-AKAP11 WT cells to sgAKAP11 knockout cells. Each waterfall plot, RIα phosphorylation sites have been annotated along with other sites of significance. **D, E.** Analysis of in vitro PKA holoenzyme activity using adapted KiMSA activity assay. Fluorescent peptide shift comparing inhibitory capacity of Flag-PKA R1α WT, phosphomimetic mutant (S83E) or phospho-null mutant (S83A), and S101A from HEK293T-sgPKA RIα lysates. 0.2uM recombinant PKA Cα, 2mM ATP and 1mM cAMP added where indicated. (E) In gel fluorescence quantification of technical triplicates of in vitro PKA kinase activity. **p = 0.0014, ***p = 0.0009. **F.** Immunoblot of co-immunoprecipitation from HEK293T-sgPKA RIα that were transiently transfected with Flag-Metap2 or Flag-PKA RIα with indicated mutations. Cells were serum starved (0.5% FBS+DMEM) for 2 hours and then stimulated with 0.5μM forskolin or DMSO for 20 minutes. **G.** Waterfall plot of statistically significant foldchanges between sgAKAP11 cells rescued with 4ALIR AKAP11 vs WT AKAP11.

By comparative phosphoproteomic analysis of the AKAP11-deleted and reconstituted samples (carried out, as in i3 neurons, under standard growth conditions), we detected 81 phosphorylated peptides that changed significantly as a function of AKAP11 status upon normalization by total levels of their respective proteins (supplementary data 3). We focused on phospho-peptides that, if increased by deletion of AKAP11, were decreased back toward control levels by re-expressing wild-type AKAP11 and, vice-versa, phospho-peptides whose abundance was decreased by AKAP11 deletion and restored by re-expressing wild-type AKAP11 (Fig. 6B-6C, supplementary data 3).

Compared to i3 neurons, fewer canonical PKA substrates were differentially phosphorylated in an AKAP11-manner in HEK-293T cells, possibly reflecting a predominant role for AKAP11 in the brain. However, in agreement with the i3 neurons one of the most hypo-phosphorylated peptides in AKAP11-deleted cells that were restored by AKAP11^WT^ re-expression was Ser83 in RIα (Fig. 6B-6C). As mentioned above, the location of S83 within the hinge-loop region of RIα suggests that its phosphorylation may regulate Cα activation. Supporting this idea, phosphorylation of nearby Ser101 by cGMP-dependent protein kinase (PKG) was shown to destabilize IS-Cα interaction, thereby contributing to PKA activation^56^.

To dissect the role of S83 phosphorylation, we carried out a modification of previously described semi-reconstituted phosphorylation assays^56,60^. A fluorescent synthetic peptide encoding for a PKA consensus phosphorylation sequence (LRRASLG) was incubated with cell lysates from RIα-deleted HEK-293T cells that were reconstituted with wild-type, phosphomimetic (S83E) or phospho-null (S83A) RIα constructs. Adding the synthetic PKA substrate peptide to the RIα-deleted lysates resulted in partial, cAMP-stimulated peptide phosphorylation by endogenous Cα present in the lysate (Fig. 6D-6E).

In lysates from cells reconstituted with wild-type RIα, reformation of the PKA holoenzyme partially inhibited Cα activity, leading to significant reduction of the phosphorylated peptide upon cAMP addition (Fig. 6D-6E). The phospho-null S83A mutant was less capable of inhibiting cAMP-induced Cα kinase activity, resulting in more phosphorylated peptide. Conversely, the phospho-mimetic S83E mutant inhibited substrate peptide phosphorylation more potently than wild-type RIα (Fig. 6D-6E). Consistent with a previous report^56^ mutating Ser 101, the target of PKG, to Ala also impaired inhibition of Cα kinase activity by RIα (Fig. 6D-6E).

To determine whether the different peptide phosphorylation efficiencies reflect different strength of association between RIα and Cα, we carried out co-immunoprecipitation experiments between FLAG-tagged, wild-type and mutant RIα, and endogenous Cα (Fig. 6F). These experiments showed that, relative to wild-type RIα, the phosphomimetic S83E RIα mutant had increased binding to Cα in cAMP-stimulated conditions, providing a rationale for the increased potency of this mutant at inhibiting Cα (Fig. 6F). In contrast, we did not detect destabilization of the RIα−Cα interaction by the S83A mutation, likely reflecting a more subtle effect of this mutation.

Because, like in i3 neurons, RIα phosphorylation at Ser83 was strongly decreased in AKAP11-depleted cells, AKAP11 may favor interaction of the bound PKA holoenzyme with one or more S83-phosphorylating upstream kinase(s) (see Discussion). Notably, Ser83 phosphorylation was significantly increased in AKAP11-deleted cells reconstituted with AKAP11^4ALIR^ mutant compared to AKAP11^WT^-reconstituted cells, even after normalizing for the total levels of RIα protein (Fig. 6G). Thus, scaffolding of the Cα−RIα-AKAP11 complex on autophagosomes may restrict its accessibility by S83-phosphorylating kinases, thereby promoting PKA activation.

The phosphoproteomic analysis in this reconstituted system identified additional AKAP11-dependent signaling events that were regulated by AKAP11 interaction with the autophagic machinery. AKAP11 deletion caused increased phosphorylation of NCK associated protein 5 like (NCKAP5L), a microtubule plus ends-binding protein that regulates microtubule bundling and acetylation^61^ at two non-PKA sites: Ser440 and Ser767. Both phosphorylation events were suppressed back to control levels by re-expressing wild-type AKAP11, but not by AKAP11^4ALIR^ (Fig. 6B, 6C and 6G). Thus, AKAP11 association to autophagosomes may be required for its ability to promote NCKAP5L dephosphorylation.

Collectively, these data point to a model where binding of AKAP11 to autophagosomes regulates its bound PKA holoenzyme in two ways: 1-by gating access to PKA by upstream RIα-phosphorylating kinases that may fine-tune the effect of local cAMP 2-by promoting autophagic degradation of the entire complex. Both processes are required for the correct phosphorylation of both PKA-dependent and independent sites, enabling precise regulation of a broad signaling program that may play important roles in neuronal cell homeostasis (Fig. 7).

**Figure 7.**
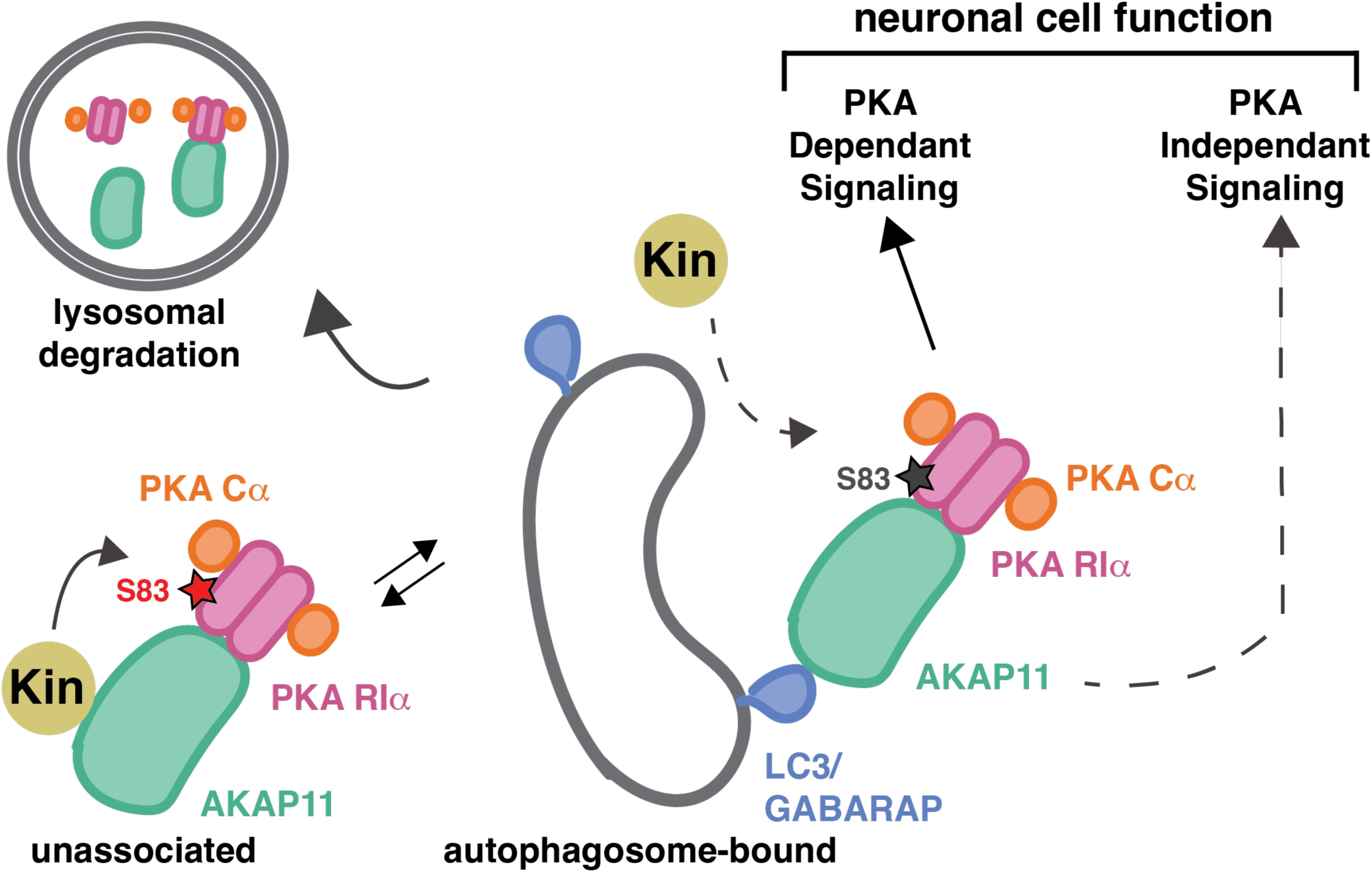
Model for AKAP11-dependent regulation of PKA signaling. The interaction of AKAP11 to LC3/GABARAP on the autophagosome is proposed to regulate association of AKAP11-bound PKA with upstream regulators, including the Ser83-phosphorylating kinase on RIα (labeled as ‘Kin’). Subsequent closure of the autophagosome and fusion with lysosomes promotes wholesale degradation of the AKAP11-bound PKA. Blocking PKA-autophagosome association leads to dysregulated PKA signaling, partly through increased RIα phosphorylation by upstream kinases. The balance between autophagosome association and degradation regulates phosphorylation of both PKA-dependent and PKA-independent substrates, thus fine-tuning downstream programs required for neuronal cell function.

## Discussion

We undertook this study with the initial goal of identifying multiprotein signaling complexes that associate with, and may be regulated by the autophagy machinery. Lysosomal immunoisolation and proteomics identified the AKAP11-bound PKA holoenzyme as a prominent kinase complex that is scaffolded to the autophagosome and is targeted for autophagic degradation. Through follow-up functional experiments and phosphoproteomics, we uncover new molecular functions of AKAP11, provide insights into how its autophagic capture modulates PKA signaling, and begin to shed light on how loss of AKAP11, an event strongly linked to the pathogenesis of SCZ/BP, impacts phosphorylation events inside neuronal cells. Our study thus uncovers a critical role for autophagosomes as physical scaffolds that govern AKAP11-PKA access to upstream regulators and downstream substrates, as well as control the overall level of this complex.

One expected function of AKAP11 is to bring the PKA holoenzyme in proximity to specific substrate proteins, facilitating their phosphorylation upon local elevation of cAMP levels and de-inhibition of bound Cα. Our phosphoproteomic results indeed support this role for AKAP11^13,16^. However, they also identify two additional functions of AKAP11. One is its ability to cause degradation of its bound RIα and Cα in autophagosomes; the second is to enable regulatory phosphorylation of its bound RIα by one or more upstream kinases, leading to stronger inhibition of Cα. The scaffolding function predicts that ablation of AKAP11 should lead to decreased phosphorylation of a subset of PKA substrates, whereas the pro-autophagic and RIα-regulating functions predict increased phosphorylation of certain PKA substrates upon AKAP11 loss (Fig. 7).

Indeed, phosphoproteomic experiments in i3 neurons show that AKAP11 loss increased phosphorylation of some canonical PKA substrates, while decreasing phosphorylation of others (Fig. 5D and supplementary data 2). Among the most hypophosphorylated PKA substrates upon AKAP11 depletion was Ser12 of RPL34; we speculate that AKAP11 may bring the RIα-Cα in proximity to RPL34 at some point during the life cycle of the ribosomal large subunit, thus favoring Ser12 phosphorylation.

Conversely, in AKAP11-defective cells the total levels of RIα and, to a lesser degree, Cα were increased, whereas the inhibitory phosphorylation of Ser83 on RIα was decreased. Combined, these effects may result in association of more numerous, overactive holoenzyme molecules with other AKAP proteins, leading to hyperphosphorylation of other PKA substrates. Indeed, depleting AKAP11 from i3 neurons led to increased phosphorylation of canonical PKA sites on multiple proteins. Thus, AKAP11 emerges as a modulator that, via its scaffolding and autophagic adaptor functions, shapes both the strength and specificity of PKA signaling toward distinct classes of substrates.

Ser83 on RIα scored as the top AKAP11-dependent phosphorylated site. Although Ser83 is not in the IS segment, it lies nearby in a flexible loop that is not resolved in the numerous published structures of RIα but can be visualized in Alphafold predictions (Fig. S3C). Though the exact molecular details of how Ser83 phosphorylation alters PKA activity is difficult to determine without structural information, the Alphafold model suggests that a phospho-group on Ser83 side chain may alter the packing of this flexible loop against PKA Cα, possibly forming a salt bridge with nearby Arg257 and thus stabilizing the interaction with the catalytic subunit.

How AKAP11 promotes RIα phosphorylation on Ser83 remains to be determined. Based on high-throughput profiling of human S/T kinase specificity^62^, Ser83 is a target site for Kinase Interacting with Stathmin (KIS), several Tyrosine kinase-like (TKL) including Activin receptor type-1B (ACVR1B) and Bone morphogenetic protein receptor type-2 (BMPR2), as well as the Casein Kinase (CK) group including CKI8 and CKIψ2. Conceivably, AKAP11 may bind to one or more of these kinases, holding them in physical proximity to the RIα linker-hinge region. Interestingly, single nucleotide polymorphisms in the KIS gene have also been associated with schizophrenia^63,64^.

The observation that, in cells expressing the AKAP11^4ALIR^ mutant, S83 phosphorylation was increased after normalizing for total RIα levels suggests that scaffolding of the Cα-RIα-AKAP11 complex to autophagosomes may regulate access of S83-phosphorylating kinases to RIα. The progressive maturation of an open phagophore into a closed autophagosome is expected to gradually restrict access of LC3-bound PKA by other interactors^26,27^. On the other hand, the confined environment within a growing phagophore could favor interactions between PKA and specific substrates, especially those that may harbor LIR motifs. It is tempting to speculate that the rate of autophagosome growth may dictate the duration of specific interactions between AKAP11-PKA and its regulators and substrates, but further studies are required to test this possibility.

Our data do not support a previously proposed model where selective autophagic degradation of RIα frees up Cα in order to amplify PKA signaling^24,43^. First, under all paradigms tested we found that a pool of cellular Cα was scaffolded to ATG8 proteins via RIα and AKAP11, and was degraded along with RIα in an autophagy- and AKAP11-dependent manner, the only exception being when we induced cAMP levels that are considered supra-physiological (Figs 1-4). This finding is in line with reports that the R- and C-subunits remain associated, at least in part, under conditions where PKA is catalytically active^46,47^. Second, ablation of AKAP11 in i3 neurons and HEK-293T cells did not uniformly lead to suppression of PKA signaling, instead it resulted in some substrates being hyperphosphorylated, while others were hypophosphorylated. (Figs 5-6).

These results indicate that AKAP11-dependent autophagic degradation of PKA is not a general ‘brake-release’ mechanism for PKA signaling. Instead, it could represent a fine-tuning mechanism that reduces the amount of PKA that is allowed to get into contact with specific substrates, as well as the duration of these PKA-substrate interactions. It should be noted that AKAP11-associated PKA activity has not been linked to specific G-protein coupled receptors (GPCRs). The localization of AKAP11-PKA in intracellular vesicles such as autophagosomes may not allow its canonical regulation through rapid cycles of local cAMP production by adenylyl cyclase followed by cAMP breakdown by phosphodiesterases^3,65^. Instead, proper regulation of this complex may rely on wholesale degradation via autophagy, coupled with phosphorylation in critical sites such as the hinge-loop region of RIα, which could reinforce or antagonize the effects of local cAMP concentrations.

It should be pointed out that in previous studies connecting autophagy to PKA regulation, no phosphoproteomic analysis of AKAP11-deficient cells was carried out. Instead, the phosphoproteomes of wild-type versus autophagy-impaired cells were compared^25,43^. Given its pleiotropic roles, autophagy controls neuronal cell homeostasis through both PKA-dependent and independent mechanisms, thus providing a rationale for the significant differences between those studies and ours. Moreover, conditions that stimulate autophagy, such as glucose withdrawal^24^ or amino acid starvation^43^ could bias AKAP11-degradation toward RIα and away from Cα, but whether and how this occurs remains to be determined.

Our phosphoproteomic analysis reveals that the signaling functions of AKAP11 go well beyond direct regulation of PKA. In the i3 neuron model, close to 95% of differentially phosphorylated peptides were non-PKA substrates. Some of these altered phosphorylation events may be indirectly linked to dysregulated PKA. However, given the ability of AKAP11 to scaffold other signaling proteins such as GSK3β and PP1, AKAP11-dependent but PKA-independent regulatory actions are likely and must be taken into account when considering how AKAP11 gene loss may alter neuronal cell homeostasis and circuitry, leading to SZ/BP. Notably, our proteomic-based lysosome profiling did not detect GSK3β or PP1 as autophagic substrates (Fig.1 and supplementary data 1); thus, another possible function of AKAP11-dependent autophagy may be to eliminate AKAP11-PKA complexes that are unassembled with the other components of this signaling supercomplex, thus helping maintain its correct composition and stoichiometry.

## Acknowledgements

The authors thank R. Irannejad for providing reagents and for critical reading of the manuscript, Z. Yue for sharing AKAP11-deleted HEK-293T cells; H. An and W. Harper for the FIP200-deleted HEK-293T cells; M. Ward for the CRISPRi-i3iPSCs containing neurogenin2 (NGN2) inducible cassette and dCas9-BFP-KRAB.

## Materials

### Antibodies and Chemicals

Reagents were obtained from the following sources: antibodies to AKAP11 (LS-C374339-100) from LSBio. FLAG (#14793), HA (#3724) PKA RI-α (D54D9) (#5675), PKA C-α (D38C6) (#5842), phospho-PKA substrate (#9624), LC3B (D11) (#3868), Vinculin (E1E9V) (#13901), TAX1BP1 (D1D5)(#13901), V5 rabbit (#13202), V5 mouse (#80076), SQSTM1/p62 (#397749), PP2AA (#2041), HOP (#5670), eEF1A (#2551), CACYBP (#3354), HSP90 (#4874), CCT2 (#3561), PSMA2 (#11864), PP2AC (#2259), GPI (#57893), GSK3*β* (#12456), Synapsin (#4297), Oct4 (#27505) from Cell Signaling Technologies. VAPA (15275-1-AP), SQSTM1/p62 (18420-1-AP), eEF1D (10630-1-AP), CCT5 (67400-1-Ig), MAP2 (17490-1-AP) from Proteintech. GFP (SC-9996), LAMP2 (SC-18822) from Santa Cruz Biotechnology. VAPB(A0302-894A), ORP11 (A304-581A) from Bethyl Laboratories. GFP (A11122) from Invitrogen, PSMD7 (ab11436) from Abcam, PKAR2A (VPA00905) from BioRad, Tuj1 from Biolegend (801202). Bafilomycin A1 (Alfa Aesar, J61835), Torin1 (Tocris, 4247), Forskolin (Cayman Chemical: 11018), Lenti-X concentrator (Takara Bio #631232). GFP-trap (gta) and mNeonGreen-trap (nta) from Proteintech/Chromotek. Pierce Anti-HA magnetic beads (88837) from Thermo Scientific. Anti-Flag beads (Sigma-Aldrich: A2220)

## Methods

### Mammalian Cell Culture

HEK293T cells and their derivatives were cultured in DMEM base media with 10% fetal bovine serum (FBS) and supplemented with 2mM Glutamine, and 1% penicillin and streptomycin. All cell lines were cultured at 37°C and 5% CO_2_. All cell lines were free of mycoplasma contamination, routinely checked by mycoplasma PCR Detection Kit (abm, #G238) and/or DAPI staining.

### Lentivirus production and infection

Lentiviruses were made by co-transfecting pLJM1 constructs with psPAX2 and pMD2G packaging plasmids into HEK293T cells using PEI transfection reagent. Viral supernatant was collected after 48 hours and again after 72 hours post-transfection and filtered using 0.45 µm syringe filter. The collected virus was concentrated using Lenti-X concentrator (Takara Bio #631232) according to manufacturer’s protocol and stored at −80°C. For lentivirus transfection, target cells were seeded along with the virus and 10ug/mL polybrene. After 24 hour incubation, virus containing media was removed and fresh media containing puromycin(1 *µ*g/mL) hygromycin (200 *µ*g/mL), or blasticidin (5 *µ*g/mL) was added for selection.

### Drug Treatments

All drug treatments were performed as follows unless otherwise specified. Bafilomycin A1 (Alfa Aesar, J61835) was used at 500 nM for 5 hours. Torin1 (Tocris, 4247) was used at 250 nM for 1 hour. Forskolin (Cayman Chemical) was used at 1 *µ*M for 25 minutes.

### Generation of CRISPR Knockout cell lines

HEK293T sgATG7 were generated as described previously (Abu-Remaileh M et al. 2017), sgFIP200 were generated as described previously (An, H. et al. 2020).

To generate HEK293T sgAKAP11, sgPRKAR1A, the following targeting sequences were cloned into pLentiCRISPRv2 vector: AKAP11-5’-ATGTCCCAGGACATTCACTG-3’, PRKAR1A-5’-ACCAAAAGATTACAAGACAA-3’. Infected cells were selected for hygromycin resistance. Cells were maintained in selection medium for 3-5 days to ensure knockout. Knockouts were validated by immunoblotting and LysoIP.

### Generation of CRISPRi i3 Neurons

To generate sgAKAP11 KD i3 Neurons, the following target sequences were cloned into pLG15 vector plasmid: sgNT 5’-GTCCACCCTTATCTAGGCTA-3’,sgAKAP11^#1^ 5’-AGCCTCCGCGGCGAGCACGT-3’, sgAKAP11^#3^ 5’-GGTGACATGTCTGTGAGCTG-3’, sgAKAP11^#5^ 5’-TCGGCGCCCGGCTCACCTGG-3’

### i^3^Neuron differentiation

CRISPRi-i^3^iPSCs containing neurogenin2 (NGN2) inducible cassette and dCas9-BFP-KRAB were a generous gift from Dr. Michael Ward (Tian et.al, 2019). CRISPRi-i^3^iPSCs were differentiated as previously described (Fernandopulle, 2018). Briefly, iPSCs were dissociated with Accustase (Gibco) and seeded onto matrigel (Corning) coated six-well plates. iPSCs were infected with lentiviruses concentrated in essential-8 medium (Gibco) expressing sgNT or sgAKAP11. After 24 hours, media was replaced with fresh media containing puromycin (1 ug/mL, SUPPLIER). Media was replaced daily with fresh media containing puromycin for 72 hrs. After selection, cells were expanded for one passage before dissociating and seeding cells into induction medium containing N2-supplement (N2,Gibco) in Knockout DMEM/F:12 with non-essential amino acids, GlutaMax and doxycycline (NEAA, Gibco; GlutaMax,Gibco; Doxycycline 2μg/mL,). Media was replaced with fresh induction media for 72 hrs, before dissociating cells and seeding onto PLO/Laminin coated plates (prepared as described in Fernandopulle, 2018) containing BrainPhys neuronal differentiation media supplemented with B27+ containing BDNF, NT-3, and GDNF. i^3^Neurons were given 60% media changes (remove 50%, add back 60%) every 3 days until harvested for experiments.

#### Lysosome immunoprecipitation (LysoIP)

Lysosomes from stable cells lines stably expressing TMEM192-RFP-3xHA were purified as previously described (Lim et al. 2019). In brief, cells were 18million cells were seeded in a 15cm dish, the following day cells were treated either with DMSO, 500nM BAFA1 for 5 hours or 20*µ*M Leupeptin + 20*µ*M Pepstatin for 24 hours. All following steps were done with cold buffers or on ice to maintain samples at 4C. Cells were washed and then resuspended in 5ml of PBS. Samples were spun at 320xg and pelleted cells were resuspended in 750uL K-PBS (136mM KCl, 10mM KH2PO4, pH7.25, with addition of fresh 0.5mM TCEP and Pierce Protease inhibitor tablet (Thermo A32965) with 3.6% Opti-prep (Sigma D1556). Cells were mechanically lysed by five passages through a 23-gauge needle and post-nuclear fractions were collected after a 1370xg 10min spin and incubated with magnetic anti-HA (Catalogue #) beads for 20min. Samples were washed 3 times with 1mL of K-PBS using a magnetic stand. For western blot analysis, samples were eluted directly into 2xUrea sample loading buffer (150 mM Tris, pH 6.5, 6 M urea, 6% SDS, 25% glycerol, 5% β-mercaptoethanol, 0.02% bromophenol blue) at room temperature overnight. For proteomics experiments, lysosomal immunoprecipitates were eluted from beads using 0.1% NP-40 in PBS for 30 min at 37°C, beads were removed and the resulting eluate was snap-frozen with LN_2_.

### LysoIP mass spectrometry and proteomic analysis

Samples were prepared in biological triplicates and analyzed by tandem mass spectrometry with an Orbitrap analyzer by the Proteomics Core Facility at the Whitehead Institute. The samples were TCA-precipitated, resuspended in TEAB, reduced and alkylated, digested, isotope-labelled, combined, cleaned up (SPE) and fractionated.

The resulting data was filtered to exclude contaminants listed in the cRAP database (common Repository of Adventitious Proteins, GPM) or in MaxQuant, and to include only proteins with peptides occurring in at least two replicates.Consequently, the dataset was normalized in Python: First, based on the mean total intensity per MS run, all MS channels were corrected for sample loading. Second, Internal Reference Scaling (IRS) was the last normalization step. Batchcorrection with Combat’s PyCombat served as benchmark. For Principal Component Analysis, the data was standard-scaled. Following normalization, the foldchange between samples was calculated as the log base 2 of the ratio of the averages. Significance for each foldchange was determined by taking the -log base ten of a welch two-tailed t-test.

### Bioinformatic analysis

List of lysosomal proteins was obtained experimentally (954 human proteins). Protein interactions between the initial list of lysosomal proteins and other interactors were determined based on BioGRID (https://thebiogrid.org/), a data base of protein-protein interactions from both high-throughput datasets and individual focused studies. The resulting list included the protein-protein interactome of the initial lysosomal protein list. The connectogram was constructed using Cytoscape (https://cytoscape.org/).

Gene ontology analysis was performed using Panther with default settings. The plot was generated in GraphPad Prism. Pathway enrichment analysis in i3 Neurons was performed using Metascape^66^. The plots were generated in GraphPad Prism.

### Immunofluorescence

Cells were seeded on fibronectin-coated glass coverslips in 6-well or 12-well plates the day before the experiment was to be performed. Cells were Bafilomycin or Torin treated where indicated and fixed using 4% paraformaldehyde (PFA) for 15 minutes at RT. Cells were rinsed 3 times with PBS, permeabilized with 0.1% (w/v) saponin in PBS for 10 minutes at RT. Cells were rinsed in PBS 3 times. Primary antibodies were diluted in 5% normal donkey serum (NDS) (Jackson ImmunoResearch) and incubated at RT for 1 hour. The coverslips were rinsed with PBS 3 times. Coverslips were then incubated in fluorescently-conjugated secondary antibodies diluted to 1:4000 diluted in 5% NDS for 45 minutes at RT while being protected from light. Coverslips were then rinsed with PBS 3 times and mounted on glass slides using VECTASHIELD Antifade Mounting Medium with or without DAPI (Vector labs)

### Microscopy

All confocal microscopy was using a spinning-disk confocal system built on a Nikon Eclipse Ti microscope (Nikon Instruments) with Andor Zyla-4.5 sCMOS camera system(Andor Technology) using a Plan Apo 60× oil objective. Images of fine cellular detail were acquired with an additional 1.5× magnifier.

### Image Analysis

For quantification of co-localization, 5–10 non-overlapping images were acquired from each coverslip. Raw, unprocessed images were imported into ImageJ v1.53 and converted to 8-bit images, and images of individual channels were thresholded independently to exclude background and non-specific staining noise and converted to binary masks. Co-localization analysis was assessed using the threshold ‘AND’ function. Percent co-localization was calculated by dividing co-localized value by lysosomal marker threshold.

### Cell lysis and Co-Immunoprecipitation

Cell lysates were prepared by removing media and rinsing cell monolayer once with DPBS and lysed in either of the following lysis buffers: Triton-based (1% Triton X-100, 10 mM sodium β-glycerol phosphate, 10mM sodium pyrophosphate, 4 mM EDTA, 40 mM HEPES, pH 7.4, and one EDTA-free protease inhibitor tablet per 50 ml) or CHAPS-based (0.3% CHAPS, 10 mM sodium β-glycerol phosphate, 10mM sodium pyrophosphate, 2 mM EDTA, 20 mM HEPES, 150 mM NaCl, pH 7.6, one EDTA-free protease inhibitor tablet per 50 ml). Cells lysed on nutator for 10 minutes. Lysates were cleared via centrifugation using a microcentrifuge at 17,000xg for 10 minutes at 4°C. Protein content in lysate samples were measured by Bradford assay or BCA. Samples of equal protein concentration and addition of 5X SDS sample buffer (235mM Tris, pH 6.8, 10% SDS, 25% Glycerol 25% *β*-mercaptoethanol, 0.1%bromophenol blue) were prepared for SDS-PAGE.

For Flag or HA immunoprecipitations, cells were seeded in a 10cm at a density appropriate for them to reach confluency after 24h. Cells were lysed as described above. 25 *µ*l of a well-mixed 50% slurry of anti-Flag M2 affinity gel or HA magnetic beads were added to each lysate sample and incubated at 4°C while on the rotator for 1 hour. For GFP or mNeonGreen IP, 15 *µ*l of well-mixed 50% slurry of GFP-Trap agarose or mNeonGreen-Trap agarose were added to each lysate sample and incubated at 4°C while on the rotator for 1 hour.

Immunoprecipitant beads were washed three times with lysis buffer. Immunoprecipitated proteins were denatured by adding 100*µ*l of sample buffer, left overnight at RT or heating to 95°C for 5 minutes.

For GFP-LC3 and GFP-GABARAP competition assay in cells, immunoprecipitation was carried out as described above except cells were lysed in 0.3% CHAPS buffer, containing the LIR peptides (500 *µ*M). Lysates were incubated at 4°C for 1.5 hours before incubating with GFP-trap agarose (10 *µ*l) for 30 minutes.

### Immunoblotting

For immunoprecipitation, 10% of the total immunoprecipitated material was loaded per lane, and 0.5% of total input was loaded per lane. Proteins were transferred to a PVDF membrane (Millipore IPVH00010), blocked with 5% non-fat milk in TBS-T, and incubated in primary antibodies (diluted in 5% milk in TBS-T) for 3 hours at RT or overnight at 4°C. Membranes were rinsed with TBS-T and incubated with horseradish peroxidase conjugated anti-rabbit or anti-mouse secondary antibodies (diluted in 5% milk in TBS-T) for 1 h at room temperature. Membranes were washed again with TBS-T and incubated with Pierce ECL Western Blotting Substrate (Thermo Scientific, 32109) before being exposed to Prometheus ProSignal Blotting Film (Genesee Scientific, 30-507L).

### Phosphoproteomic sample preparation

HEK293T sgNT or sgAKAP11 and sgAKAP11 stably expressing V5-AKAP11 WT or V5-AKAP11 dLIR were seeded (5.0 x10^6^ cells) in 10cm dish in DMEM complete medium (DMEM, 10% FBS, 2mM Glutamine, 1% penicillin/streptomycin). After overnight adherence cells were switched to DMEM + 10% dFBS medium. Cells were harvested 24 hours later and lysed as previously described. Protein content in lysate samples were measured by BCA. Lysis was flash frozen in liquid nitrogen until mass spectrometry analysis.

### Phosophoproteomic Mass spectrometry

All samples were labeled with iodoacetamide, and resuspended in 100mM of 4-(2-Hydroxyethyl)-1-piperazinepropanesulfonic acid (EPPS) buffer, pH 8.5 and digested at 37C with trypsin overnight. The samples were labeled with TMT Pro and, quenched with hydroxylamine. All the samples were combined and desalted using a 100mg Sep-Pak cartridge, followed by drying in a rotary evaporator. Phosphopeptides were enriched using a Fe-NTA spin column (Thermo Fisher). Flow through samples were dried, and fractionated with basic pH reversed phase (BPRP) high-performance liquid chromatography (HPLC) as described previously. Samples were desalted via StageTip and dried with speedvac. Samples were resuspended in 5% formic acid, and 5% acetonitrile for LC-MS/MS analysis. Mass spectrometry data were collected using Exploris 480 or Orbitrap Eclipse mass spectrometers (Thermo Fisher Scientific) coupled with a nLC-1200 or Vanquish Neo liquid chromatograph (Thermo Fisher Scientific), respectively, with a 90 or 150 min gradient across a Nano capillary column (100µm D) packed with ∼35cm of Accucore C18 resin (Thermo Fisher Scientific). A FAIMSPro (Thermo Fisher Scientific) was utilized for field asymmetric waveform ion mobility spectrometry (FAIMS) ion separations. The instrument methods, which are embedded in the RAW files, included Orbitrap MS1 scans (resolution of 120000; mass range 400-1600 m/z; automatic gain control (AGC) target 4×10^5^, max injection time of 50 ms. MS2 scan parameters were set as described previously (CID collision energy 35%; AGC target 7.5 × 10^3^; rapid scan mode; max injection time (50mM)).

All samples were labeled with iodoacetamide, and resuspended in 100mM of 4-(2-Hydroxyethyl)-1-piperazinepropanesulfonic acid (EPPS) buffer, pH 8.5 and digested at 37C with trypsin overnight. The samples were labeled with TMT Pro and, quenched with hydroxylamine. All the samples were combined and desalted using a 100mg Sep-Pak cartridge, followed by drying in a rotary evaporator. Phosphopeptides were enriched using a Fe-NTA spin column (Thermo Fisher). Flow through samples were dried, and fractionated with basic pH reversed phase (BPRP) high-performance liquid chromatography (HPLC) as described previously. Samples were desalted via StageTip and dried with speedvac. Samples were resuspended in 5% formic acid, and 5% acetonitrile for LC-MS/MS analysis. Mass spectrometry data were collected using Exploris 480 or Orbitrap Eclipse mass spectrometers (Thermo Fisher Scientific) coupled with a nLC-1200 or Vanquish Neo liquid chromatograph (Thermo Fisher Scientific), respectively, with a 90 or 150 min gradient across a Nano capillary column (100µm D) packed with ∼35cm of Accucore C18 resin (Thermo Fisher Scientific). A FAIMSPro (Thermo Fisher Scientific) was utilized for field asymmetric waveform ion mobility spectrometry (FAIMS) ion separations. The instrument methods, which are embedded in the RAW files, included Orbitrap MS1 scans (resolution of 120000; mass range 400-1600 m/z; automatic gain control (AGC) target 4×10^5^, max injection time of 50 ms. MS2 scan parameters were set as described previously (CID collision energy 35%; AGC target 7.5 x 10^3^; rapid scan mode; max injection time (50mM)).

### Phosphoproteomics Analysis

Comparative phospho-peptide analysis between genotypes was conducted by first normalizing each phospho-peptide to account for differences in total protein abundance in each genotype. As such, the relative abundance of each protein at the whole cell level in each genotype was compared to the levels in sgNT control samples then this normalized abundance was used to normalize the corresponding phosphopeptide for each replicate. Following normalization, the foldchange between samples was calculated as the log base 2 of the ratio of the averages. Significance for each foldchange was determined by performing a welch two-tailed t-test. Phosphopeptides foldchanges with pvalues < 0.05 were then plotted on a waterfall plot. All analysis was done using custom R scripts.

### Statistical analysis

All graphs were assembled, and statistics were performed using Prism 10 (GraphPad). Error bars on all graphs are shown as the mean ± SEM. The details of each statistical test performed are given in the legend accompanying each figure. Unless otherwise indicated, all co-localization analysis was performed on 10 non-overlapping fields that contain a minimum of 3 cells per field. Unless otherwise indicated, all proteomic measurements were performed on three independent biological replicates for each sample.

### Kinase Activity Assay

HEK293T-sgPKA RIa cells were seeded in 6cm tissue culture-treated plates. After overnight incubation, cells were transiently transfected with 5*µ*g Flag-RIa WT type and mutant DNA using PEI, as described previously. 24 hours post transfection, cells were lysed in the plate using 120ul of cold NDP-40 Lysis buffer (1X PBS, 1% NDP-40, 1X PhosStop, 1X EDTA-free protease inhibitor). Lysates were cleared via centrifugation using a microcentrifuge at 17,000xg for 10 minutes at 4°C and then normalized to 2*µ*g/*µ*L. Each kinase reaction was made by using a previously described protocol with final concentrations of 200mM Tris-HCl pH 7.4, 10mM MgCl_2_, 0.2mM ATP, 0.5mM TCEP, 1X EDTA-Free protease inhibitor, 1X PhosStop, 21uM 5’FAM-Kemptide (Anaspec) and 0.4 *µ*g/*µ*L cell lysate. 1mM cAMP was added where noted. Kinase reactions were incubated at RT for 20 minutes, followed by the addition of 5X SDS sample buffer (235mM Tris, pH 6.8, 10% SDS, 25% Glycerol 25% *β*-mercaptoethanol, 0.1% bromophenol blue). 2.5uL of each sample were run on 12% NuPAGE Bis-Tris protein gels with MES running buffer. Blank lanes were left in between each sample lane and no protein ladder standard was used.

In-gel fluorescence was imaged using ChemiDoc. Fiji was used for quantification of fluorescence. Background subtraction was performed using a 50-pixel rolling ball subtraction followed by intensity measurements for each individual band. Fraction of substrate phosphorylated was calculated as phosphorylated intensity/(phosphorylated intensity + unphosphorylated intensity).

---

**Supplementary Figure 1.**
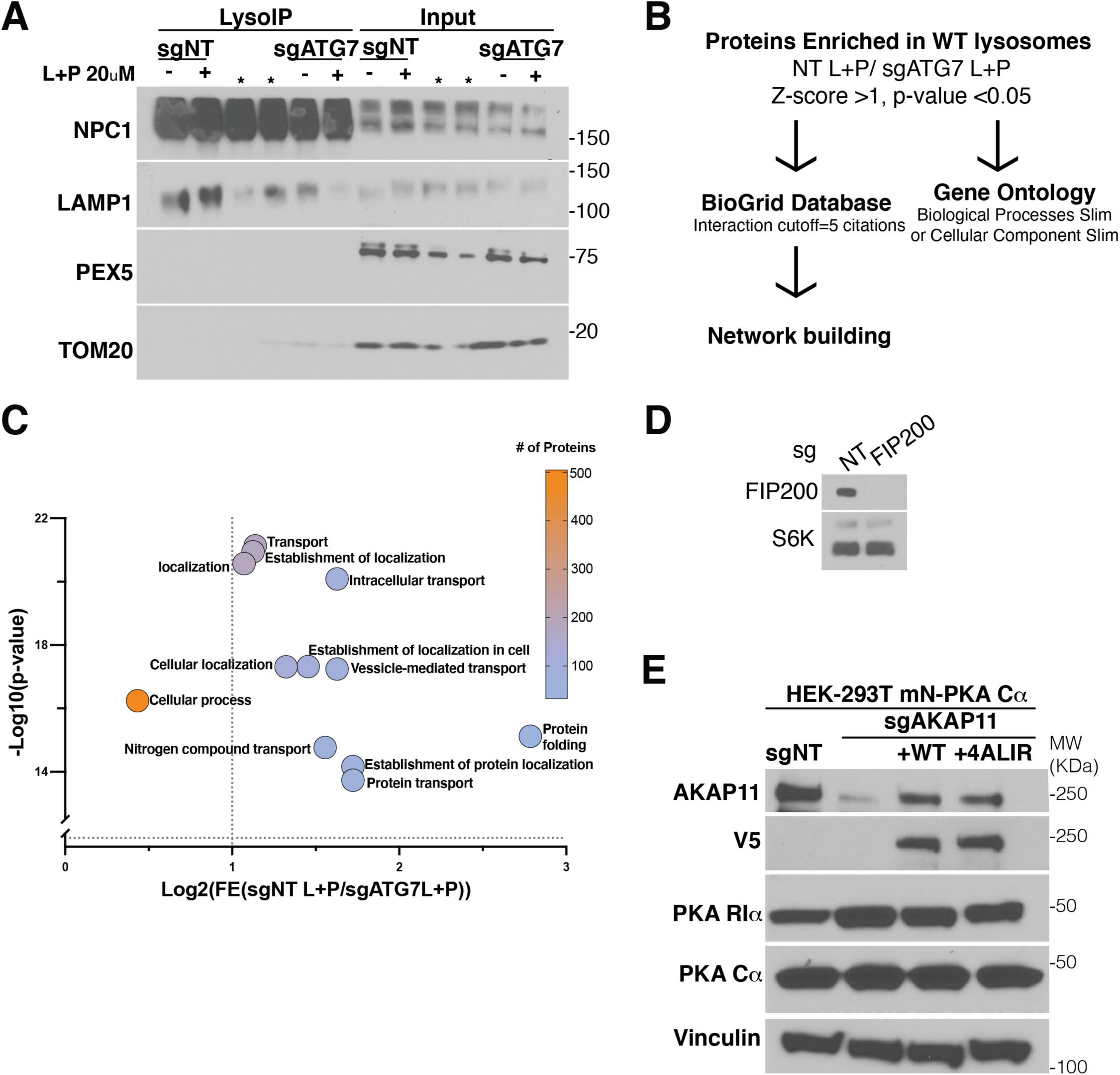
**A.** Immunoblots of Lysosomal immunoprecipitation and corresponding input from HEK-293T-sgNT or sgATG7 after treatment 20*µ*M Leupeptin and 20*µ*M Pepstatin for 24 hours to block lysosomal degradation of lysosomal substrates. **B.** Outline depicting bioinformatic pipeline in which proteins identified as ‘Hits’ (Log2(FC[sgNT L+P/sgATG7 L+P])>1, p-value<0.05) in LysoIP from WT and ATG7-null cells were subjected to custom-written pipeline where interactors of Hits (using citation cutoff of ≥ 5 citations) were identified using data from BioGrid. List of protein ‘Hits’ were also entered in panther to generate a gene ontology analysis for enrichment of biological processes or enrichment of cellular component. **C.** Volcano plot of “biological processes slim” Go-terms enriched in wild-type lysosomes compared to autophagy-null lysosomes in 20*µ*M L+P 24-hour treatment. **D.** Western blot validation of FIP200 knockout in HEK293T endogenously tagged with mNeon-PKA Cα cells as compared with sgNT cells. **E.** Immunoblot validation of AKAP11 knockout using whole cell lysate from HEK293T sgNT or sgAKAP11 and sgAKAP11 with stably expressing the indicated V5-AKAP11 construct.

**Supplementary Figure 2.**
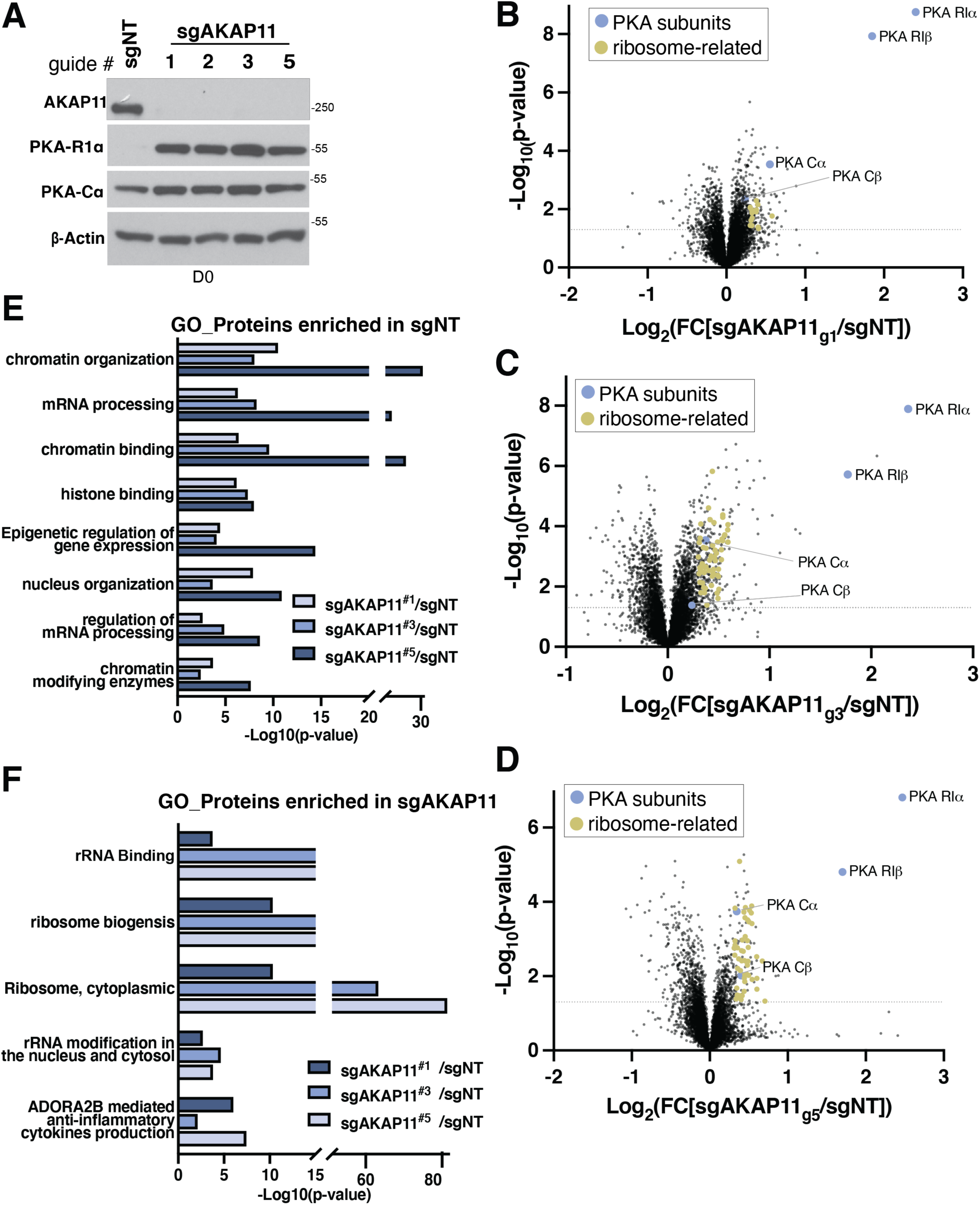
**A.** Immunoblot of whole cell lysate of DIV0 i3 neurons validating degree of knockdown of four sgAKAP11 guides compared to sgNT. **B, C, D.** Proteomic analysis of DIV7 i3 neurons in 3 different guides targeting AKAP11. Blue nodes indicate PKA subunits. Gold nodes indicate ribosome-associated proteins (ribosome, cytoplasmic and ribosome biogenesis). **E, F** Pathway enrichment analysis of DIV7 i3 neurons comparing (E) pathways enriched in sgNT (F) pathways enriched in sgAKAP11. Proteins included in the pathway enrichment analysis had a −0.2≥Log_2_FC≥ 0.2 and p-value <0.05.

**Supplementary Figure 3.**
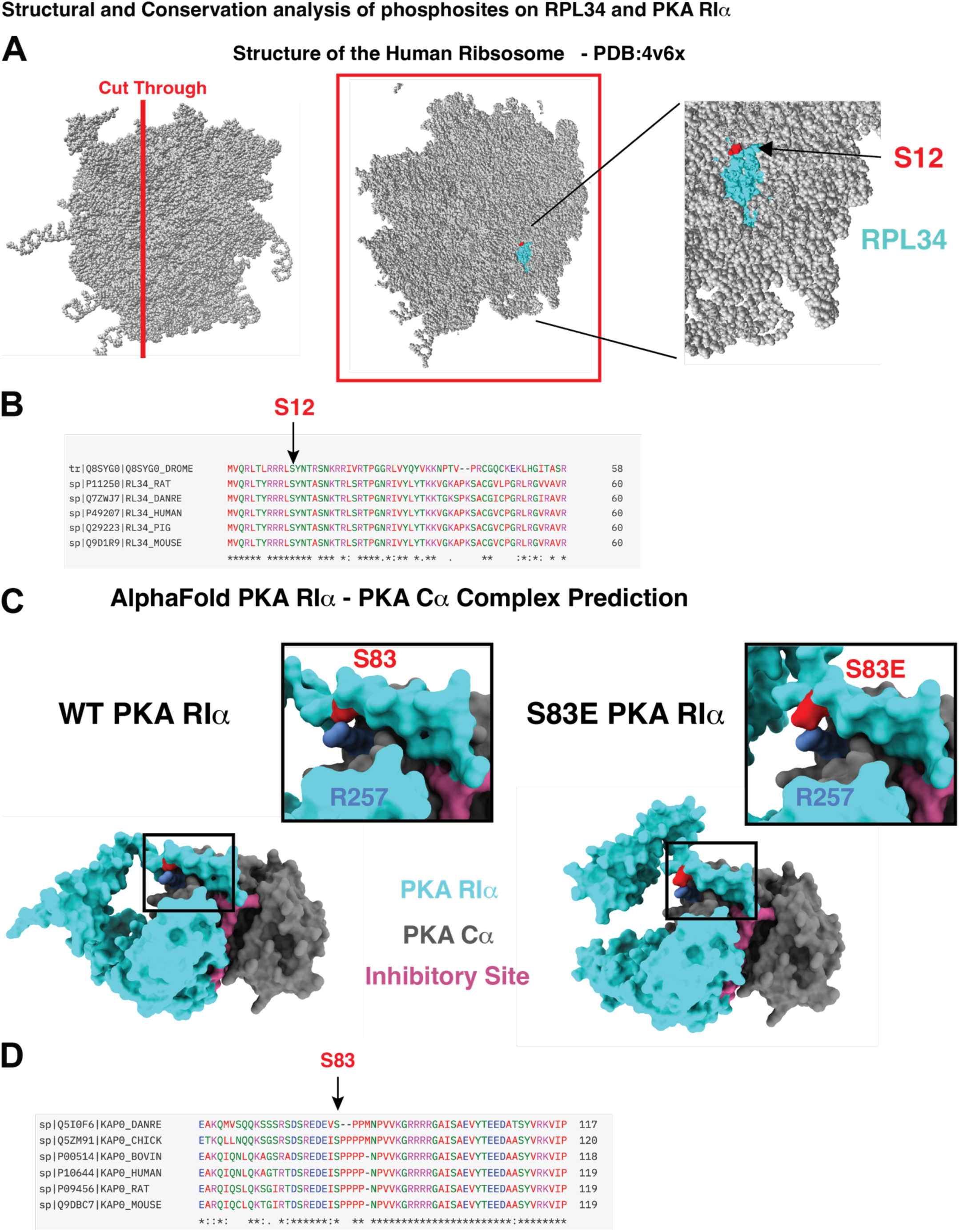
Structural and Conservation analysis of phosphosites on RPL34 and PKA RIα. **A**. The cryoEM structure of the human ribosome with RPL34 (cyan) deeply buried in the complex and Ser12 annotated in red. **B**. Sequence alignment of RPL34 across species showing conservations of Ser12. **C**. Alphafold prediction of the full length PKA holoenzyme complex, both WT and containing S83E mutant RIα. PKA RIα contains an inhibitory site (pink) that inserts in the active site of PKA Cα (gray). An unstructured loop nearby contains Ser83 (red). The S83E mutation, which mimicks phosphorylation, is predicted to form a salt bridge with Arg257 of Cα and induce a significant conformational rearrangement in RIα. **D**. Sequence alignment of PKA RIα across species showing conservation of Ser83.

**Supplementary Figure 4.**
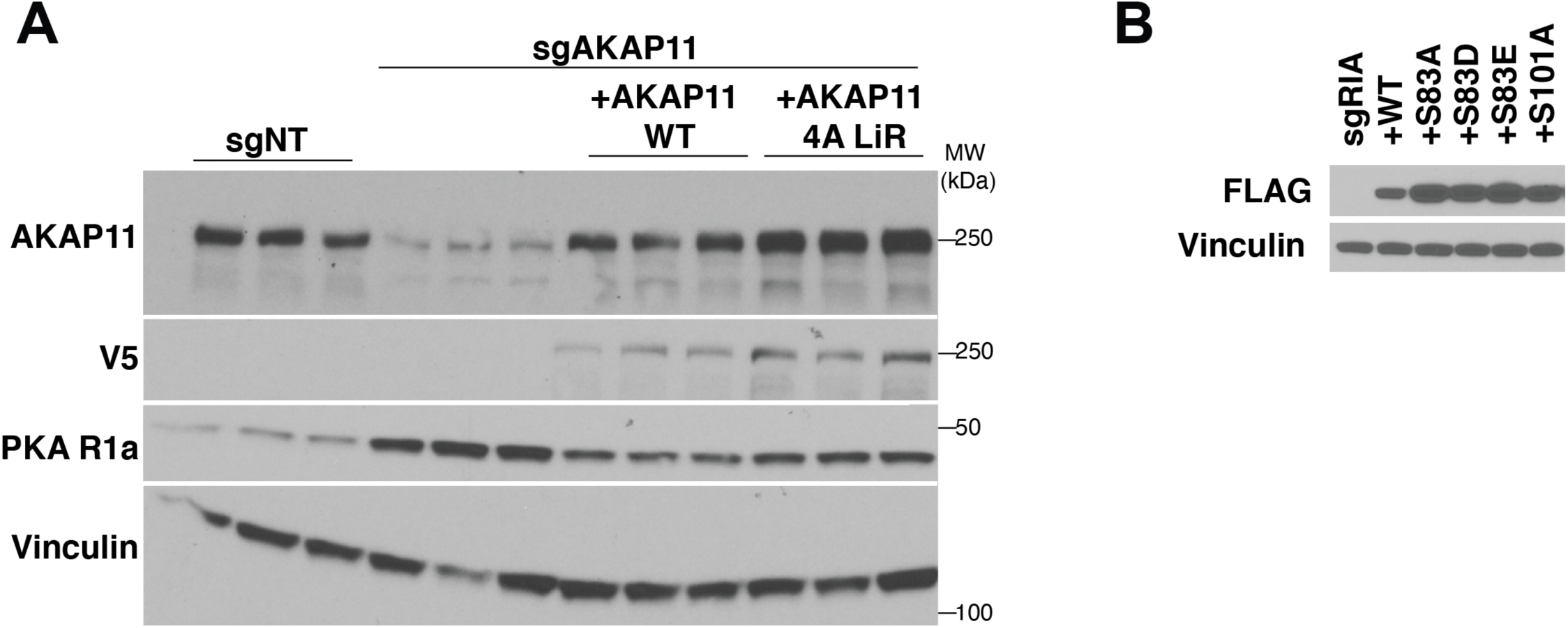
**A.** Immunoblot of whole cell lysate from HEK293T sgNT and sgAKAP11 and sgAKAP11 cells stably expressing the indicated V5-AKAP11 construct in triplicate. Cells were grown in DMEM+10% dFBS. These samples were used for phosphoproteomic experiments. **B.** Western blots of sgRIα lysates used in Kemptide kinase assay showing vinculin as a loading control and transient expression of FLAG-RIα rescue constructs.

